# Can hubs of the human connectome be identified consistently with diffusion MRI?

**DOI:** 10.1101/2022.12.21.521366

**Authors:** Mehul Gajwani, Stuart J. Oldham, James C. Pang, Aurina Arnatkevičiūtė, Jeggan Tiego, Mark A. Bellgrove, Alex Fornito

## Abstract

Recent years have seen a surge in the use of diffusion MRI to map connectomes in humans, paralleled by a similar increase in processing and analysis choices. Yet these different steps and their effects are rarely compared systematically. Here, in a healthy young adult population (n=294), we characterized the impact of a range of analysis pipelines on one widely studied property of the human connectome; its degree distribution. We evaluated the effects of 40 pipelines (comparing common choices of parcellation, streamline seeding, tractography algorithm, and streamline propagation constraint) and 44 group-representative connectome reconstruction schemes on highly connected hub regions. We found that hub location is highly variable between pipelines. The choice of parcellation has a major influence on hub architecture, and hub connectivity is highly correlated with regional surface area in most of the assessed pipelines (ρ>0.70 in 69% of the pipelines), particularly when using weighted networks. Overall, our results demonstrate the need for prudent decision-making when processing diffusion MRI data, and for carefully considering how different processing choices can influence connectome organization.

**Author Summary:** The increasing use of diffusion MRI for mapping white matter connectivity has been matched by a similar increase in the number of ways to process the diffusion data. Here, we assess how diffusion processing affects hubs across 1760 pipeline variations. Many processing pipelines do not show a high concentration of connectivity within hubs. When present, hub location and distribution vary based on processing choices. The choice of probabilistic or deterministic tractography has a major impact on hub location and strength. Finally, node strength in weighted networks can correlate highly with node size. Overall, our results illustrate the need for prudent decision-making when processing and interpreting diffusion MRI data.

**Code and data availability:** All the data used in this study is openly available on Figshare at https://doi.org/10.26180/c.6352886.v1. Scripts to analyze these data are available on GitHub at https://github.com/BMHLab/DegreeVariability.

**Competing Interests:** The authors declare that they have no competing interests.

## Introduction

A major priority for neuroscience is to robustly and accurately map the connections of the human brain (Sporns et al., 2005). These connections are thought to be distributed heterogeneously across different brain regions, with putative ‘hub’ areas having stronger and more frequent connections with other regions (Arnatkevičiūtė et al., 2021; Oldham et al., 2019; van den Heuvel & Sporns, 2013b). Hub connectivity is viewed as playing an integral role in supporting coordinated dynamics (Mišić et al., 2015; van den Heuvel & Sporns, 2013a). It has a distinct developmental trajectory (Fan et al., 2011; Oldham et al., 2022; Oldham & Fornito, 2019); is important for cognitive function (Fagerholm et al., 2015; Sleurs et al., 2021); is under strong genetic influence (Arnatkevic□iūtė et al., 2018, 2019, 2021; Fulcher & Fornito, 2016); and is implicated in a diverse range of clinical disorders (Crossley et al., 2014; de Lange et al., 2019; Fornito et al., 2015; Gollo et al., 2018). In humans, the anatomical connectivity of hub and non-hub brain regions is most commonly mapped using tractographic analysis of diffusion magnetic resonance imaging (MRI) data (Betzel & Bassett, 2017; Sotiropoulos & Zalesky, 2019). One challenge of such analyses is that diffusion MRI data are noisy and the final generation of a tractographic estimate of connectivity – a tractogram – depends on many different processing steps, each relying on multiple user-selected options (Jones & Cercignani, 2010; Oldham et al., 2020; Sarwar et al., 2021). As a result, different investigators make different choices, resulting in connectome models that arise from data processed in different ways.

Amongst the numerous available processing choices, commonly varied steps include: the algorithms used to seed, propagate, and prune tractography streamlines in individuals (Jeurissen et al., 2019; Sarwar et al., 2019); the cortical parcellations used to delineate distinct regions and facilitate computational tractability (Lawrence et al., 2021); and the approaches used to generate a group-representative network (Betzel et al., 2019). One recent preliminary investigation found that changing preprocessing steps can shift the location of hubs from parietal/cingulate cortex to temporal cortex (Oldham et al., 2020). Other work has similarly found a high variability in the validity of streamline reconstruction between research groups utilizing different processing pipelines (Maier-Hein et al., 2017). Valid inferences about the structure and function of human connectome hubs critically depend on our ability to reliably identify them, but a detailed examination of precisely how variations in connectome-generation pipelines affect classifications of network hubs has not yet been conducted. Here, we evaluate how such variations influence hub identification, focusing on three key steps in the connectome generation pipeline: tractography algorithm, cortical parcellation, and group reconstruction.

Tractography refers to the process by which white matter streamlines are generated based on anisotropic water diffusivity. An indirect marker of connectivity with many distinct steps (Jeurissen et al., 2019), it is dependent on user-defined parameters that include (amongst others) where the streamlines are seeded, how they propagate, and where they can terminate. One well-known, significant choice is between probabilistic and deterministic tractography. In some instances, probabilistic tractography has been shown to match more closely with *ex vivo* anatomical tract dissections than deterministic tractography (Lilja et al., 2014), while other work has reported that probabilistic tractography is more prone to generating false positive connections (Sarwar et al., 2019). Indeed, there is a general trade-off between the sensitivity and specificity of different tractography algorithms, with probabilistic tractography being more sensitive but less specific compared to deterministic tractography (Thomas et al., 2014). Moreover, the use of a particular tractography algorithm may interact with other choices in diffusion MRI pipelines, further contributing to connectome variability. For instance, Li et al. (2012) found that 50% of hubs are re-categorized when streamline seeds (the locations from where streamlines are propagated) are located at the grey matter-white matter interface rather than deep in the white matter. Methods to differentially retain anatomically probable streamlines have also been suggested (Schiavi et al., 2020; Smith et al., 2012), and manual inclusion/exclusion of streamlines for a given bundle have been shown to increase reconstruction accuracy from 73% to 91% compared to template-generated dissections (Schilling et al., 2020). As such, changing the parameters used for generating tractograms in individuals can result in connectomes with significantly different architecture, a phenomenon which has not been extensively characterized.

Cortical parcellations – the atlases used to define the boundaries between brain areas acting as network nodes – are also a source of variability in connectome architecture. Such parcellations have been undergoing continual revision since at least the time of Brodmann (Brodmann, 1909; Zilles, 2018) and the methods used to generate them are highly variable (Arslan et al., 2018). For example, parcellations have been generated using manual segmentation based on sulcal and gyral anatomy (Desikan et al., 2006); using network models based on functional connectivity (Schaefer et al., 2018); and on multimodal combinations of anatomical, microstructural, and functional features (Genon et al., 2018; Glasser et al., 2016; Wang et al., 2015). Parcellations are also difficult to compare due to differences in the number of regions delineated (Fornito et al., 2010; Zalesky et al., 2010), variability in the surface areas of regions (Van Essen et al., 2012), and inter-hemispheric (a)symmetry (Yan et al., 2022).

After the connectomes of individuals have been constructed, it is common practice to aggregate the data to derive a group-representative network (de Reus & van den Heuvel, 2013a; Yeh et al., 2016). At this stage it is important to define which connections (edges) should be maintained and how these connections should be weighted. Different methods have been proposed for the former, including retaining edges that are the strongest or most frequently occurring across individuals (de Reus & van den Heuvel, 2013a), retaining edges that are the least variable across people (Roberts et al., 2017), or retaining edges that preserve a specific proportion of connections in different distance bins (Betzel et al., 2019). Complicating the methodological differences of these approaches, the specific thresholds used are often chosen heuristically (Bordier et al., 2017), making it difficult to compare studies using different thresholds.

Here, we compare the effects of different choices at these three key steps –– tractography, parcellation, and group reconstruction –– on properties of hub connectivity in a sample of 294 healthy young adults. The different options examined at each step resulted in 1760 different group-representative connectomes. We evaluate the effects of each of these choices on measures of binary and weighted node degree, given that these measures are fundamental to many other network measures and to the definition of network hubs. In particular, we focus on both the distribution of degree measures across nodes and their spatial topography, evaluating the consistency with which hubs are localized to the same anatomical positions.

## 1. Results

The processing steps that we independently varied are summarized in Figure 1. In total, we compared 1760 distinct pipelines. This encompassed 40 pipelines for generating individuals’ connectomes (10 tractography workflows and 4 parcellation schemes) as well as 44 pipelines for reconstructing group-representative connectomes (4 group aggregation methods and 11 density thresholds). The Results section is organized as follows: first, we examine the effects of different processing steps on statistical properties of the weighted degree distributions. Second, we compare the spatial distribution of node degrees across the different pipelines. Finally, we examine how specific properties of each parcellation are associated with node degree. We focus in the main text on analyses of weighted node degree (also called node strength) distributions and report results for unweighted (binary) distributions in the Supplementary Material.

**Figure 1:**
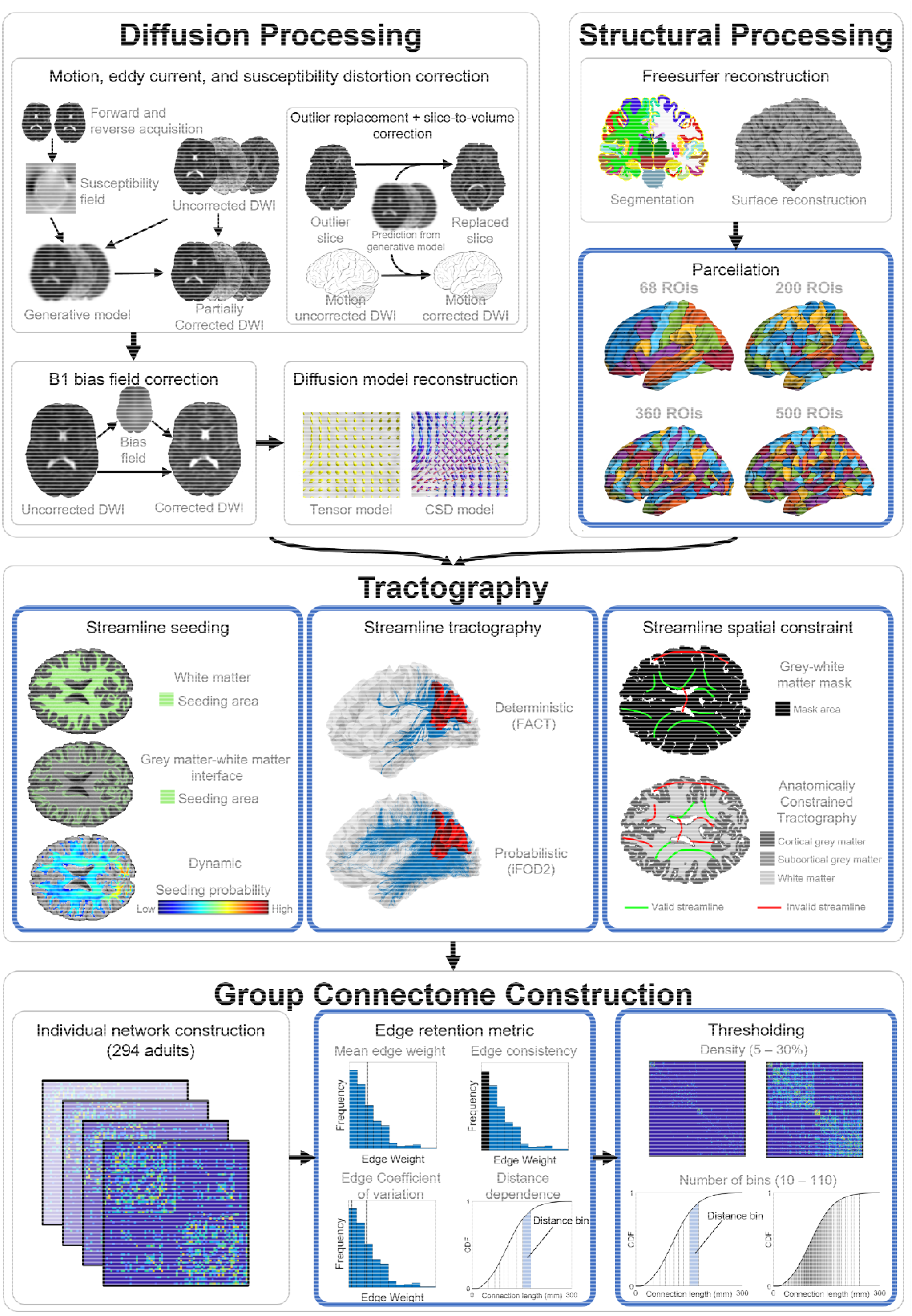
Workflow of processing steps used in group connectome construction. Outer gray boxes group multiple options were compared. The first three boxes (diffusion processing, structural processing, and tractography) refer to the reconstruction of streamlines within one individual. The fourth box (group connectome construction) refers to the process by which the connectome matrices of individuals are used to generate the group representative connectome. Note that structural processing is also used to inform individual network reconstruction. DWI = Diffusion Weighted Imaging; ROIs = Regions of Interest; FACT = Fibre Assignment by Continuous Tractography; iFOD2 = second-order Integration over Fibre Orientation Distributions. Adapted from (Oldham et al., 2020), licensed under CC-BY-4.0.

### 1.1. Statistical properties of node degree distributions

Figure 2 shows how properties of the node strength distributions vary as a function of parcellation and tractography parameters. For simplicity, we focus on networks thresholded at a connection density of 20% and aggregated using edge CV, since different density thresholds and aggregation methods did not substantially alter the shape of the distribution (Figures S1-4). We focus on three key properties that quantify the tail decay of the empirical degree distributions in comparison to the exponential distribution: the right-tailedness (Jordanova & Petkova, 2017), the skewness, and the excess kurtosis (see Meth-ods for details). The exponential distribution has been generally defined as the cutoff for heavy-tailed distributions (Foss et al., 2013), and is therefore a benchmark for assessing whether the concentration of connectivity in hub nodes exceeds that of a single-scale network (Amaral et al., 2000). The distribution of outliers (the right-tailedness) and the third and fourth standardized moments (the skewness and kurtosis) have been previously described to capture tail behavior in statistical dis-tributions (DeCarlo, 1997; Jordanova & Petkova, 2017; Westfall, 2014). They are also parameter-invariant for the exponential distribution (right-tailedness ≈ 0.009, skewness = 2, and excess kurtosis = 6), allowing for assessment of heavy-tailedness unbiased by user-defined parameters.

**Figure 2:**
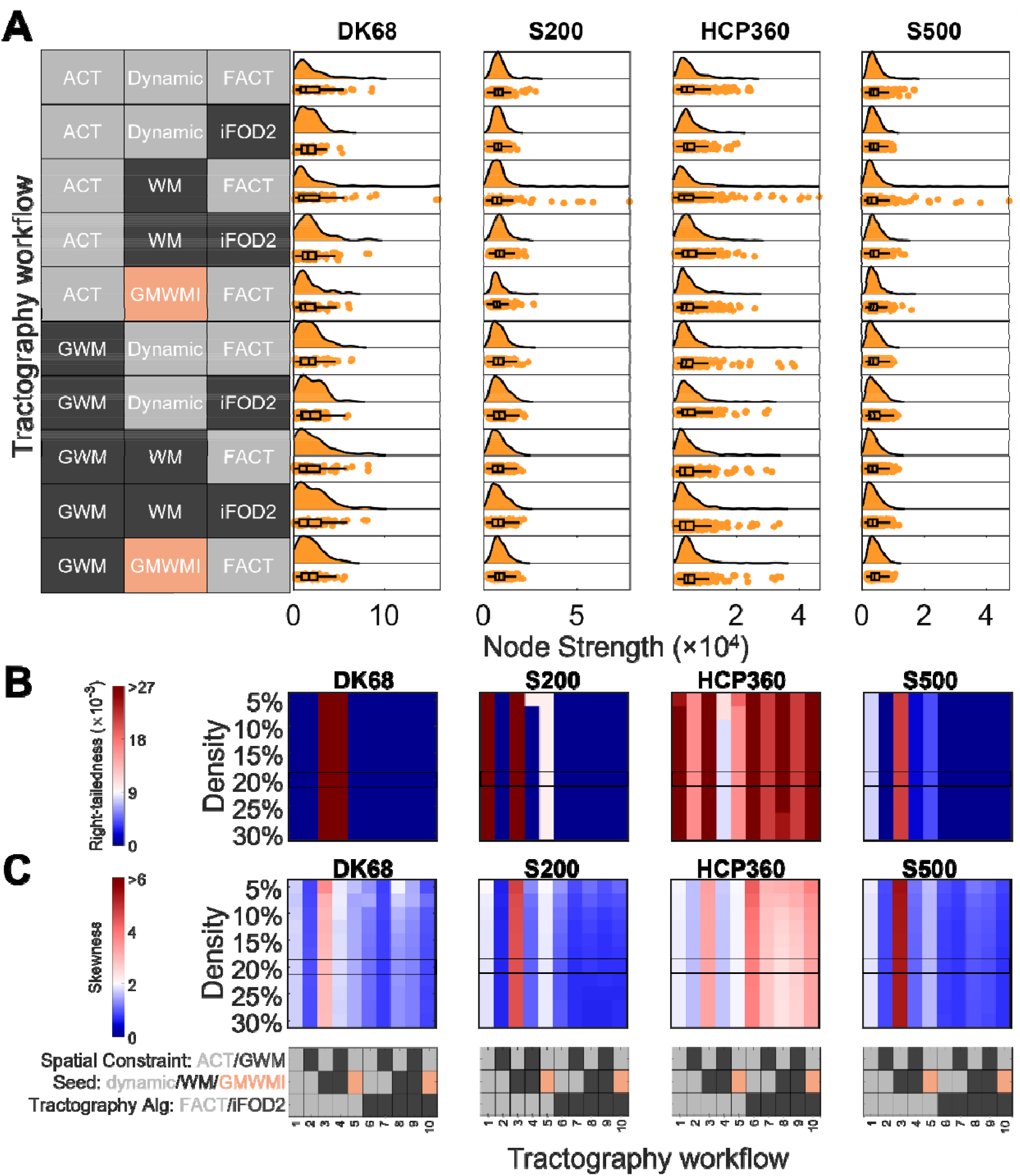
Effect of processing on weighted connectome strength distributions. (A) Node strength distributions from each of the 10 tractography workflows and 4 parcellation schemes assessed. Here, the group connectome is reconstructed using the edge coefficient-of-variation (CV) and a connection density of 20%. (B, C) Right-tailedness (B) and skewness (C) of strength distributions in each parcellation as a function of tractography and density threshold. Here, the group connectome is reconstructed using edge CV, and the boxed row corresponds to the data in (A). The range of cool/warm colors correspond to a skewness and right-tailedness less than/greater than those of the exponential distribution. The grey/black/peach key represents the processing options used in each workflow, with the possible options for that step color-coded; further details in section 3.4. Parcellation: DK68 = Desikan-Killiany 68 nodes, S200 = Schaefer 200 nodes, HCP360 = Glasser 360 nodes, S500 = Schaefer 500 nodes. Tractography: ACT = anatomically constrained tractography, GWM = grey-white masking; Seed = streamline seeding algorithm, dynamic = dynamic seeding, WM = white matter seeding, GMWMI = grey matter-white matter interface seeding; FACT = fiber assignment by continuous tractography, iFOD2 = second-order integration over fiber orientation distributions.

Figure 2A indicates that all pipeline variations qualitatively show some evidence of skewness and a heavy tail. The DK68 and HCP360 parcellations show the largest positive skews, whereas the S200 and S500 parcellations show much smaller tails, consistent with a lower likelihood of finding very highly connected hubs. The exception is the use of Workflow 3 (ACT/WM seeding/FACT), which shows an extended tail across all parcellations.

The right-tailedness and skewness of the node strength distributions are shown in Figures 2B and 2C, respectively, as functions of parcellation, workflow, and density. 692 of 1760 pipelines (39%) have more right-sided outliers than an equivalent exponential degree distribution (Figure S2). The skewness is always positive (Figure S3), ranging between 0.42 and 6.04. However, only 25% of pipelines (432 of 1760) demonstrate a greater skew than the exponential distribution (skewness = 2). Excess kurtosis (Figure S4) is greater than 0 (i.e., more kurtotic than the Gaussian distribution) in 94% of the pipelines (1654 of 1760) and greater than 6 in 25% of the pipelines (445 of 1760). Thus, despite the widely-held belief that connectomes contain network hubs, a property that should be reflected in a heavy-tailed degree distribution, only ∼25% to ∼40% of the processing pipelines examined here displayed distributions with properties that align with this hypothesis, depending on how heavy-tailedness is quantified.

There are three further key findings that are evident in Figures 2B-C. First, tractography algorithm has a major effect on the properties of the node strength distribution, with evidence of a skewed, heavy-tailed distribution only obtained when using specific processing steps in combination with deterministic tractography (FACT). More specifically, the most skewed distributions are observed when combining FACT with ACT (workflows 1, 3, and 5), with the additional use of white matter seeding yielding the highest skewness (workflow 3). This effect is apparent across connection densities and persists regardless of the method used for group aggregation (Figures S2-4). In contrast, probabilistic tractography (iFOD2) only yields evidence of a right-tailed distribution when combined with the HCP360 parcellation.

The second key finding in Figures 2B-C is that parcellation type affects the strength distributions. The skewness, kurtosis, and right-tailedness of connectomes using the HCP360 parcellation are generally higher than other parcellations, regardless of the processing steps used. In part, this may be due to the known size discrepancy between regions of the HCP parcellation, which we examine in more detail in Section 1.3. The skewness, kurtosis, and right-tailedness only exceed those of an exponential distribution when using the HCP360 parcellation for all pipelines using probabilistic tractography.

The third key finding in Figures 2B-C is that skewness and right-tailedness change minimally as connection density is varied. Thus, connection density does not have a large impact on the tails of the strength distributions of weighted connectomes.

The results for binarized connectomes show some differences relative to weighted connectomes (Figures S5-8). Specifically, the skewness, right-tailedness, and kurtosis of binarized connectomes are more stable than weighted connectomes when different data processing parameters are varied. Only 2.1% of connectomes (37 of 1760) are more skewed than the exponential distribution (all using the HCP360 parcellation; Figure S6). Similarly, 2.2% (39 of 1760) are more kurtotic (Figure S7) and 6.2% (109 of 1760; Figure S8) show evidence of greater right-tailedness than the exponential distribution. Notably, the skewness of the binarized connectomes was more sensitive to changes in connection density, particularly when edge consistency and CV-based thresholding were used with the HCP360 parcellation. In these specific cases, the distributions showed supra-exponential skewness and right-tailedness at thresholds of 5–10% but not at thresholds of 20–30%. Evidence of strong skewness, kurtosis, or right-tailedness in connectomes using parcellations other than HCP360 was weak and only occurred in rare instances.

Our analysis of strength distributions indicates that conclusions about the degree to which connectivity is concentrated in network hubs can vary substantially depending on how the data are processed, with tractography algorithm (i.e., deterministic or probabilistic) and parcellation type having particularly large impacts. We now turn our attention to how different processing choices affect the spatial embedding of degree; i.e., we evaluate whether different pipelines produce network hubs localized to consistent anatomical regions.

For each parcellation separately, we first calculated the partial rank correlations between the degree distribution of each pair of pipelines, controlling for regional surface area. The resulting matrices (one for each parcellation) represent the similarity in spatial location of hubness between tractography workflows. Hierarchical agglomerative clustering of these matrices was used to group similar pipelines together (Figures 3A-D). Taking the S200 parcellation as an example, Figure 3B shows that there are substantial differences in the node strength correlation between pairs of workflows, spanning the range -0.11 < *ρ* < 1.00, with an average of 0.47 (Figure 3E). As per prior work (Oldham et al., 2020), two large clusters are evident, separating workflows using deterministic tractography from those using probabilistic tractography. The average correlation within the cluster corresponding to deterministic tractography is 0.64 (0.19 < *ρ* < 1.00) and is 0.67 within the probabilistic tractography cluster (0.32 < *ρ* < 0.99), with the average correlation between clusters being 0.30 (−0.11 < *ρ* < 0.57). Within the deterministic tractography cluster, there is a further split as a function of spatial constraint (i.e., ACT versus GWM) with further subdivisions according to seeding strategy. Within the probabilistic tractography cluster, smaller clusters can also be defined as a function of spatial constraint and seeding strategy, but these sub-clusters are less homogeneous than those in the deterministic tractography cluster. The basic cluster structure was largely consistent across parcellations, with some minor variations. For instance, with the DK68 atlas, connectomes generated using dynamic seeding, probabilistic tractography, and a grey-white mask (workflow 7) formed their own sub-cluster. The group aggregation algorithm and threshold density have minimal impact on the clustering (Figure S9).

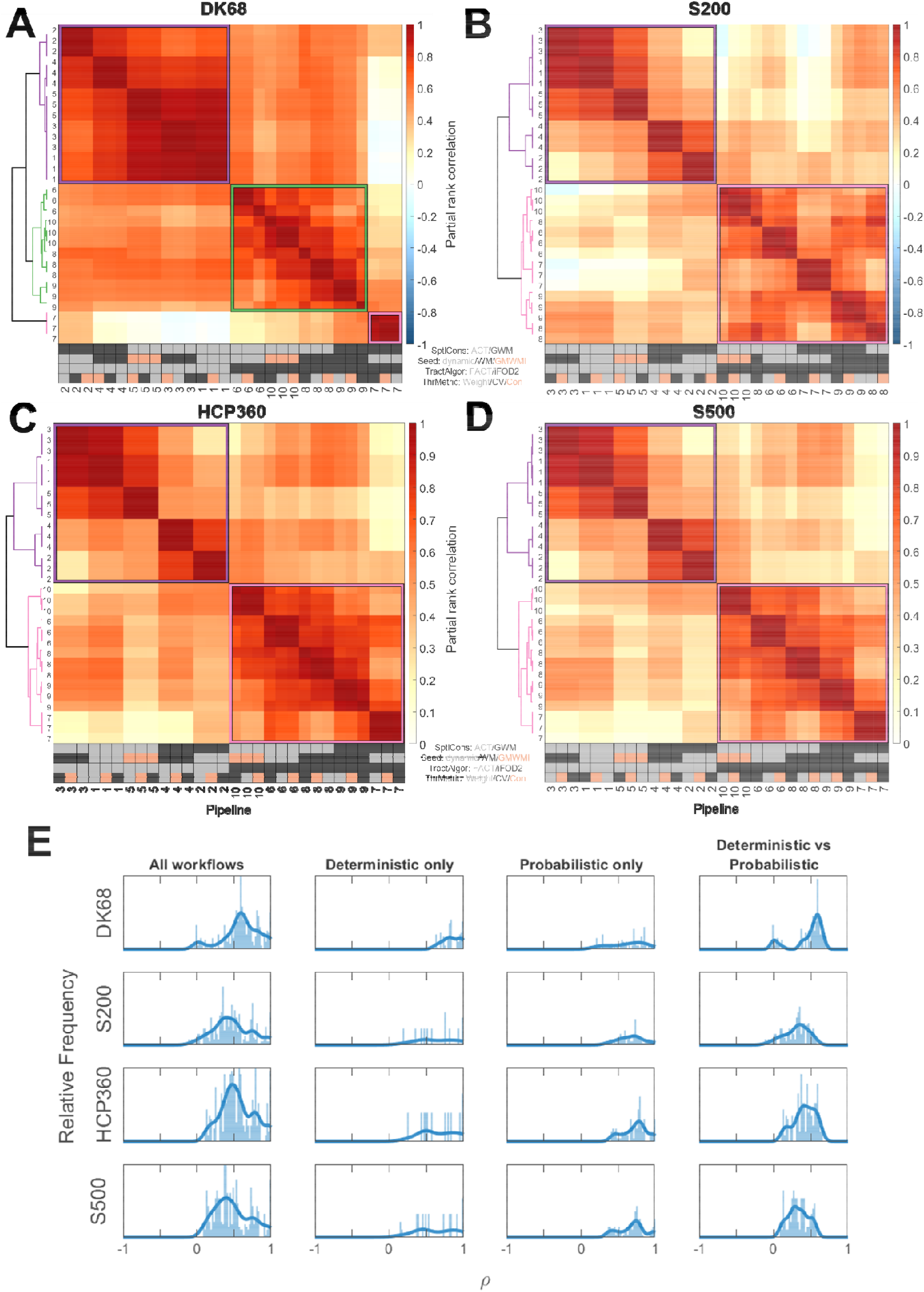
distributions between tractography workflows and group reconstruction metrics in each parcellation (A) DK68, (B) S200, (C) HCP360, (D) S500, with density 20%. Each heatmap shows partial rank correlations, corrected for surface area. Pipelines are reordered using hierarchical clustering. Pipeline numbers refer to tractography parameters; each pipeline occurs three times as three density-matched group reconstruction thresholding metrics are compared. The grey/black/peach key represents the processing options used in each workflow, with the possible options for that step color-coded; further details in section 3.4. (E) Distribution of correlation coefficients within each heatmap. Each row represents one parcellation. The first column shows the frequency of correlation coefficients across each heatmap. The subsequent columns show the subset of correlation coefficients when comparing deterministic pipelines only (second column), probabilistic pipelines only (third column), and deterministic versus probabilistic only (fourth column). Parcellation: DK68 = Desikan-Killiany 68 nodes, S200 = Schaefer 200 nodes, HCP360 = Glasser 360 nodes, S500 = Schaefer 500 nodes. Tractography: SptlCons = spatial constraints on streamline propagation, ACT = anatomically constrained tractography, GWM = grey-white masking; Seed = streamline seeding algorithm, dynamic = dynamic seeding, WM = white matter seeding, GMWMI = grey matter-white matter interface seeding; TractAlgor = Streamline tractography algorithm, FACT = fiber assignment by continuous tractography, iFOD2 = second-order integration over fiber orientation distributions. Group aggregation: ThrMetric = group-reconstruction thresholding metric, Weight = edge weight, CV = edge coefficient-of-variation, Con = edge consistency.

Figure 4 shows how the spatial distribution of node strengths varies across workflows and parcellations. First, for a fixed parcellation (e.g., the S200 parcellation), the location of putative hubs varies considerably across maps under different processing variations. When using deterministic tractography (FACT), the highest strength nodes are located in the vicinity of the paracentral lobule and supplementary motor area, compared to be located in primary visual areas when using probabilistic tractography (iFOD2). The enhanced skewness associated with the combination of ACT/WM/FACT (workflow 3) is also apparent in these maps. Notably, the DK68 and HCP360 atlas appear more robust to processing variations, which may be driven by the large variability in the size of the parcels comprising these atlases. We consider this issue in more detail in the next section.

**Figure 4:**
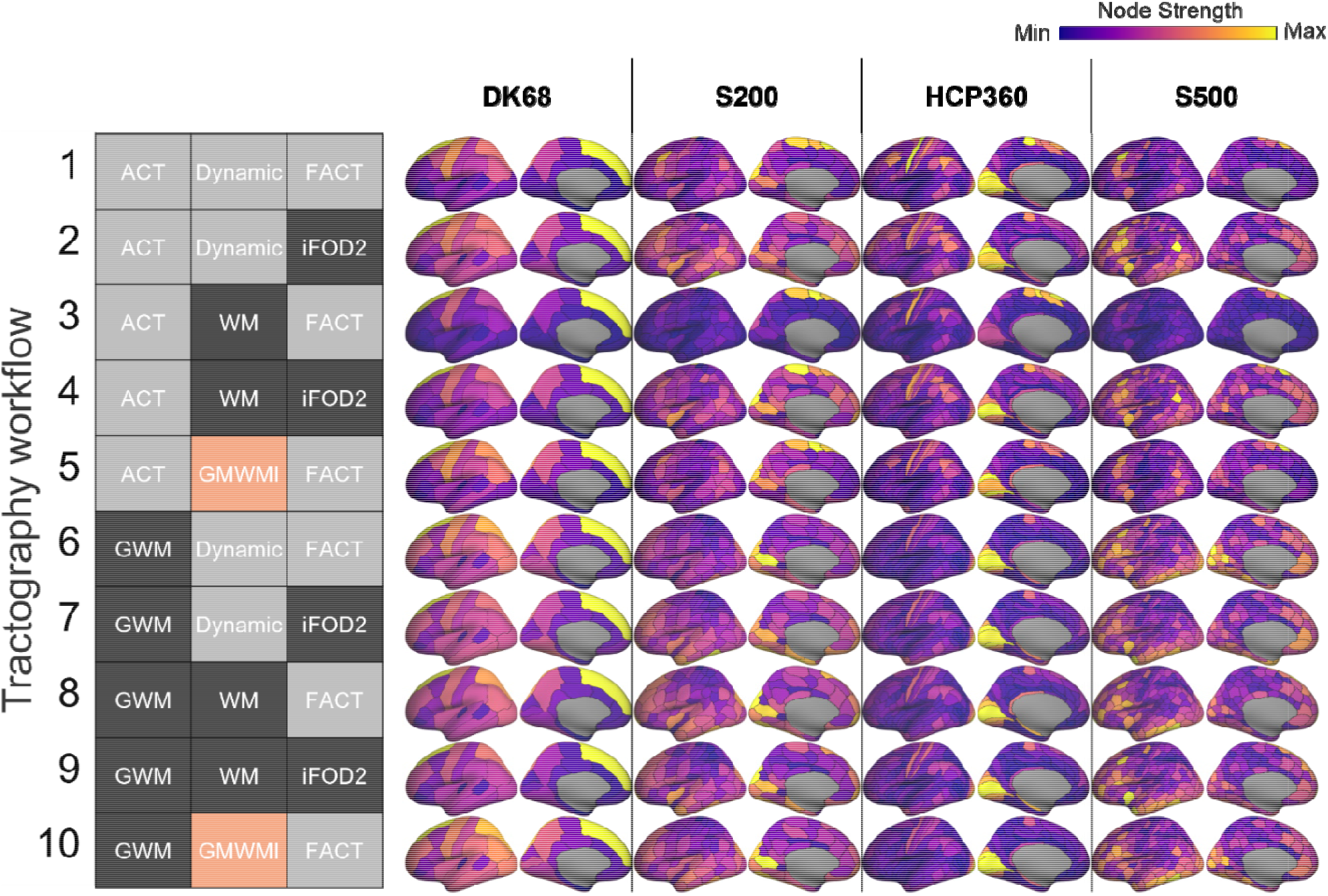
Spatial maps of node strength for each cortical parcellation and tractography workflow. The colormap is scaled independently for each image for visual purposes. Group reconstructions use edge coefficient of variation (CV) and a density of 20%. The grey/black/peach key represents the processing options used in each workflow, with the possible options for that step color-coded; further details in section 3.4. Parcellation: DK68 = Desikan-Killiany 68 nodes, S200 = Schaefer 200 nodes, HCP360 = Glasser 360 nodes, S500 = Schaefer 500 nodes. Tractography: SptlCons = spatial constraints on streamline propagation, ACT = anatomically constrained tractography, GWM = grey-white masking; Seed = streamline seeding algorithm, dynamic = dynamic seeding, WM = white matter seeding, GMWMI = grey matter-white matter interface seeding; TractAlgor = Streamline tractography algorithm, FACT = fiber assignment by continuous tractography, iFOD2 = second-order integration over fiber orientation distributions.

Second, for a fixed workflow, Figure 4 shows variations across parcellations. Such comparisons across parcellations can only be performed qualitatively as the lack of region-to-region correspondence precludes direct comparison. Once again, conclusions about the locations of hub regions vary dramatically. The highest strength nodes for the DK68 atlas are located in the medial prefrontal cortex (PFC), whereas this area is associated with relatively low strength in the other parcellations. The S200, HCP360, and S500 parcellations show a greater degree of consistency, with higher strength nodes located in visual, lateral prefrontal, anterior insula, and inferior parietal regions. The major discrepancy between these parcellations is in the primary sensorimotor cortex, which has a high strength in HCP360, but not in S200 or S500 parcellations. For a given parcellation and tractography workflow, the group aggregation algorithm and threshold density have a small effect on node strength rankings (Figure S10).

Figures S11 and S12 compare the spatial distribution of node rank variability across tractography workflows, group reconstructions, and densities. As tractography workflow is altered, relative node rankings can vary drastically (Figure S11). Whilst the middle nodes are the most variable, the rankings of the top 10-20% of nodes are also highly changeable. For example, across 30 density matched group-reconstructed connectomes, the node that is ranked 4th on average can vary between ranks 1 to 43; similarly, the node that is ranked 10th on average can vary between ranks 2 to 54. We also assessed the node rank variability across variations in group reconstruction for an exemplar tractography workflow (Figure S12). While the findings are qualitatively similar to those in Figure S11, the magnitude of the variability is much lower in this analysis. When considered together with Figure 3, this suggests that tractography workflow – rather than group reconstruction – drives much of the variance in node rankings.

Similarities between node degree distributions in binarized connectomes are shown in Figures S13 and S14. The results show a major difference between probabilistic and deterministic tractography across all parcellations (Figure S13). The locations of the strongest nodes are similarly variable: for example, using the S200 parcellation, the highest degree node is consistently found in the insula, but other high-degree nodes are located in the occipital cortex when using FACT and in temporal areas when using iFOD2 (Figure S14).

The effects of parcellation on node strength seem, in some cases at least, related to the node surface area (here, node strength distributions were observed for the DK68 and HCP360 parcellations, which have a much wider variance in regional surface areas than the S200 and S500 parcellations (Figure S15). Moreover, the medial PFC in the DK68 parcellation falls under the superior_frontal_gyrus anatomical label, which is largest region in this parcellation. In the other parcellations, the medial PFC is sub-divided into smaller parcels. It is also evident from Figure 4 that the degree sequences of the DK68 and HCP360 atlases are fairly robust to processing variations, which is notable since these are the atlases with the greatest variance in regional surface area. Areas with larger surface area will be able to accommodate more incoming and outgoing connections, and we should thus expect node strength/degree to be related with surface area. This raises the possibility that node degree will largely be driven by regional size variations, particularly in atlases with a high variance of parcel surface area. We therefore examine the degree to which the size of a node in a given parcellation determines its hubness by correlating node strength with surface area across parcellations, workflows, and group reconstruction methods.

### 1.3. The effect of variations in regional surface area

Figure 5A show spatial maps of node strengths obtained for two example parcellation and workflow combinations (S200 + GWM/dynamic seeding/iFOD2 and HCP360 + ACT/GMWMI/FACT) and Figure 5B shows the scatterplot of the association between node surface area and strength for each. Figure 5C shows the correlation coefficients for all tractography parameters and threshold densities for the S200 and HCP360 parcellations using edge CV (all tractograms/parcellations/group reconstructions are in Figure S16 and S17). Across all processing and parcellation combinations, the correlations between node strength and node surface area spanned the range 0.10 < *r* < 0.96, with a median of 0.82. Correlations for pipelines using probabilistic tractography (iFOD2) were all above r = 0.78 with a median correlation coefficient of 0.88. This high correlation persists regardless of thresholding algorithm or connection density (Figure S16). Correlations for pipelines using deterministic tractography (FACT) were somewhat lower, with a median value of 0.67 (0.10 < *r* < 0.91). The relationship between node strength and surface area was slightly weaker when using either of the Schaefer parcellations (S200 or S500) or the combination of ACT/WM/FACT (or both; Figure S16 and S17). Note that while node strength is highly correlated with node surface area, the same is not true of individual edges: Figure S18 shows that the weight of individual edges is not related to the total surface area of their endpoint nodes.

**Figure 5:**
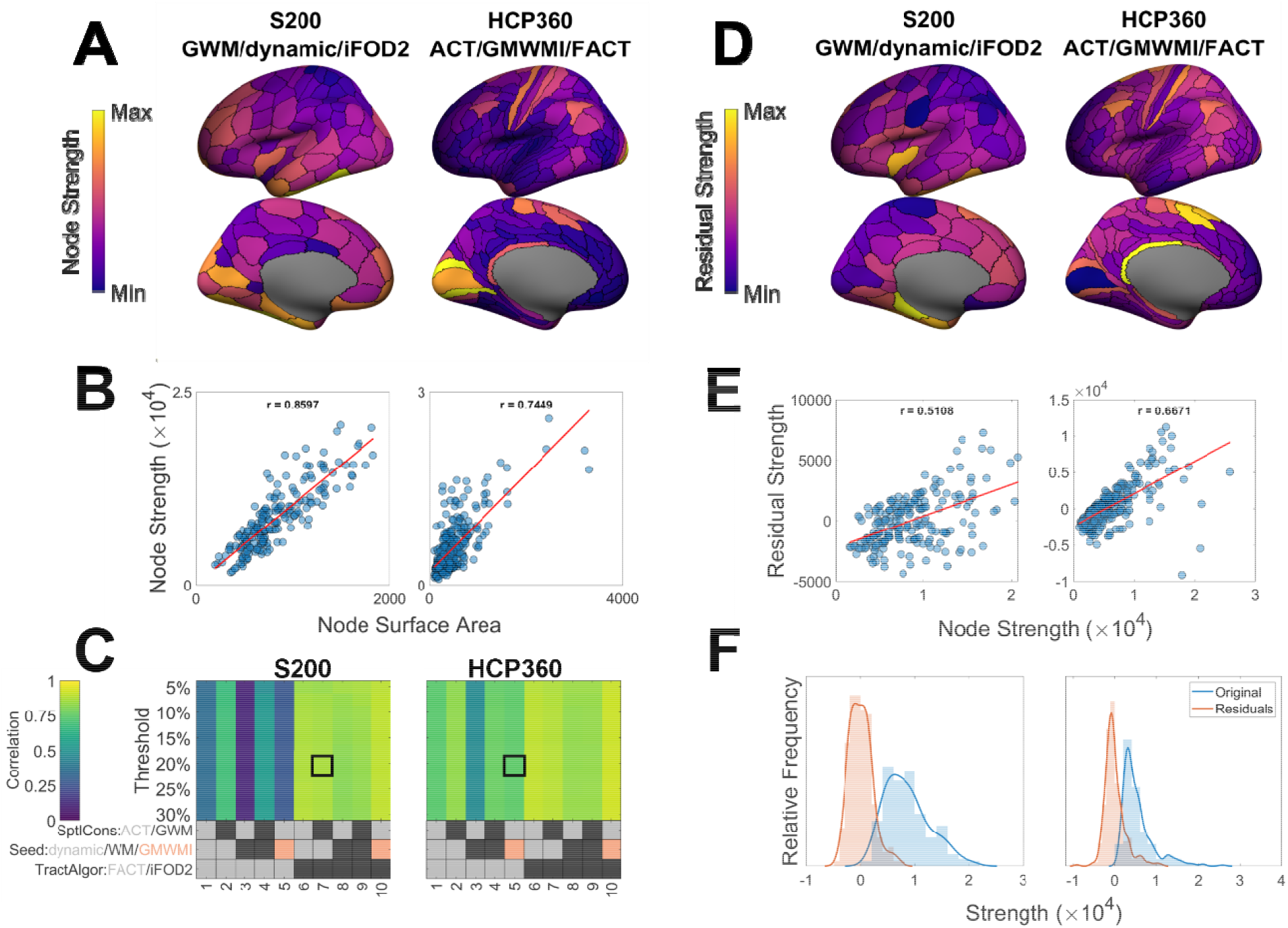
Relationship between node strength and node surface area. (A) Spatial maps of node strengths in two example parcellations/tractography workflows. In this example, connection density is 20% with group connectomes constructed using the edge coefficient-of-variation (CV). For ease of visualization, only the left hemisphere is shown. (B) Relationship between node strength and node surface area for all nodes shown in panel A. (C) Pearson’s correlation coefficient between node strength and node surface area as a function of tractography workflow and density threshold. The outlined areas (boxes) correspond to the plots in panel B. The grey/black/peach key represents the processing options used in each workflow, with the possible options for that step color-coded; further details in section 3.4. (D) Spatial maps of residual strengths when the linear relationship in panel B is removed. (E) Relationship between residual strengths shown in panel D and original strengths shown in panel A. (F) Frequency distribution of residual strengths shown in panel D and original strengths shown in panel A. Parcellation: S200 = Schaefer 200 nodes, HCP360 = Glasser 360 nodes. Tractography: SptlCons = spatial constraints on streamline propagation, ACT = anatomically constrained tractography, GWM = grey-white masking; Seed = streamline seeding algorithm, dynamic = dynamic seeding, WM = white matter seeding, GMWMI = grey matter-white matter interface seeding; TractAlgor = Streamline tractography algorithm, FACT = fiber assignment by continuous tractography, iFOD2 = second-order integration over fiber orientation distributions.

We next investigated whether removing the dependence of node strength on size changes the spatial distribution of the former measure. Figure 5D shows an example of the spatial distributions of the residual node strength values obtained after removing their dependence on regional surface area via linear regression. In the S200 parcellation, the nodes with the highest residuals tend to be those that are originally of medium-high strength (e.g., insula and inferior temporal gyrus). Thus, the locations of the most strongly connected nodes remain approximately similar. In contrast, in the HCP360 parcellation, the retrosplenial and pre-supplementary motor area cortices show disproportionately high strengths relative to surface area.

The relationship between the residuals and the original strength of each node is shown in Figure 5E. The residuals remain highly correlated with the original strengths (mean correlation across all pipelines r = 0.63 ± 0.20). Whilst the distribution of residuals may change in location (mean) and scale (variance), the skewness, right-tailedness, and kurtosis are preserved (Figure 5F). Qualitatively similar results were obtained for all parcellations and group reconstructions (Figure S19).

Figure S20 shows the relationship between node surface area and degree in binarized connectomes. Similar to weighted node strength, the correlation is stronger when using probabilistic than deterministic tractography. In contrast with node strength, binary node degree generally has a lower correlation with surface area but a greater dependence on threshold density than in weighted connectomes (Figure S20). Across all parcellations and workflows, the median correlation was 0.44 (compared to 0.82 for the weighted connectomes). However, this relationship weakened as connection density increased. For example, in the Schaefer parcellations (S200 and S500), a correlation coefficient above 0.5 occurred only when the density was below 20%. Taken together, these findings suggest that atlas-specific variations in parcel size can influence, but not fully explain, statistical and topographical properties of node strength and degree.

## 2. Discussion

We characterized the effects of several key processing steps of diffusion MRI on the distribution and location of the most strongly connected regions of the human connectome. In total, we examined 1760 group connectomes (40 pipelines for individual connectome construction, and 44 group reconstruction schemes) which represent common choices and techniques in diffusion MRI processing. However, this analysis still encompasses only a fraction of the flexibility and variability that is possible in diffusion processing pipelines.

We found that, across all the investigated pipelines, evidence of concentrated connectivity in hubs (i.e., degree distribution properties that differ from the exponential case) was apparent in only a minor fraction of pipeline variations. When relying on node strength to define hubs, variations in tractography algorithm and parcellation had a much greater effect than changes in group reconstruction method and connection density. The use of binary degree yielded a less pronounced concentration of connectivity in network hubs and the resulting connectomes are more sensitive to connection density. When considering the spatial topography of hubs, the choice between probabilistic and deterministic tractography resulted in the largest difference and, in some circumstances, led to anti-correlated weighted degree sequences. Finally, although hubs were often the regions with the largest surface area, particularly in weighted connectomes, removal of this dependence of degree on region size generally retained a similar hub topography. Together, these findings raise concerns about the consistency with which hubs can be identified in the literature and suggest that careful consideration must be paid to processing choices when mapping connectomes with diffusion MRI.

### 2.1. The effects of tractography algorithm

Degree distribution properties and hub strengths showed significant variations based on the tractography parameters used. Amongst the properties compared in our analysis, the choice of probabilistic versus deterministic tractography was shown to drive the greatest variation in degree distribution properties, as represented in the skewness, kurtosis, and right-tailedness of the degree distributions. In general, deterministic tractography resulted in more asymmetric distributions with heavier tails; in particular, the most skewed distributions in weighted connectomes resulted from the combination of white matter seeding, an anatomical streamline constraint, and deterministic tractography (ACT/WM/FACT). Given that these results were not consistently replicated across other workflows, the results of this combination of parameters may be atypical. Whether this atypicality reflects a unique sensitivity of this workflow combination in recovering the true underlying network architecture, or a result of interaction between processing steps, is unclear.

Changes in the shape of the degree distribution were also reflected in changes in the location of the strongest nodes and the relationship to node surface area. Probabilistic tractography showed a strong correlation between node strength and node surface area in weighted connectomes. This was observed across all parcellations, seeding strategies, spatial constraints, and group reconstructions. As such, the locations of hubs derived from probabilistic tractography were slightly more consistent, and degree distributions were generally more correlated between workflows.

### 2.2 The effects of cortical parcellation

Many different parcellations have been used in the literature to map connectomes. These parcellations vary with respect to two key factors relevant to connectome mapping: their spatial resolution and their variance in parcel sizes. Spatial resolution naturally affects the precision with which the connectivity of regions can be resolved and can lead to differences in the spatial topography of hubs. For instance, the medial PFC was a prominent hub in the DK68 atlas but not in the other parcellations, where this area is sub-divided into smaller regions. This variation is likely related to regional variations in surface area, since the medial PFC is among the largest in the DK68 parcellation. Such variations can interact with other processing choices; for instance, degree distributions were highly skewed and kurtotic (sub-exponential decay) when using probabilistic, but not deterministic, tractography with the HCP360 parcellation, for which the largest parcel is more than 1.5× larger than the largest parcel in the S200 parcellation.

### 2.3 The effects of regional variation in surface area

To the extent that a given parcellation defines valid functional areas of the brain, the correlation between region size and degree may be an accurate reflection of biological reality––some regions may be more connected simply because of their size. However, it can be useful to determine whether a region’s hubness is simply a result of its surface area. It is somewhat reassuring that the relative degree rankings of different areas only changed moderately after controlling for the effects of size variations, but these effects should nonetheless be considered when drawing conclusions about the hub status of specific brain regions. Further work could consider the mechanisms by which the weight of individual edges (which are uncorrelated with node surface area) contribute to total node strength (which is often highly correlated with node surface area).

### 2.4. The effects of group reconstruction and connection density

The specific method for aggregating individual connectomes into a group-averaged representation had minimal effect on node strength distributions or topographies. Binary degree was more susceptible to the effect of varying connectome density, which is likely because thresholding removes the weakest connections. Such connections make a small contribution to weighted degree but make an equal contribution to strong edges when estimating binary degree.

### 2.5. Limitations

We intentionally used model-free quantities to characterize network degree distributions to simplify and standardize measures across the various pipelines considered. An alternative is to fit specific distributions to the data. For example, previous studies have reported that weighted connectomes have a degree distribution that follows a power-law distribution (Varshney et al., 2011), a truncated power-law distribution (Modha & Singh, 2010), or a generalized Pareto distribution (Zucca et al., 2019). In the best case, these models can suggest a biological mechanism which may produce observed patterns of hub connectivity, but care should be taken in performing inference using such analyses (Clauset et al., 2009). Our approach offers a hypothesis-free way of quantifying the degree to which connectivity is concentrated in putative hub nodes, but future work could consider characterizing the precise forms of connectome degree distributions in more detail.

We focused here on cortex for simplicity, given the large number of processing pipelines that we examined. However, our conclusions are sufficiently general that they should not be significantly altered by the inclusion of non-cortical areas. Similarly, we focus on a normative cohort, given that applications to some clinical cohorts may require further steps and thereby exacerbate workflow variability (Martínez-Heras et al., 2015; Mochizuki et al., 2023; Richards et al., 2021; Sanvito et al., 2020; Shu et al., 2011). Future work may examine the effects of such additional variations (Gonzalez-Aquines et al., 2019; Horbruegger et al., 2019; Lipp et al., 2020).

The absence of a ground truth for diffusion MRI makes comparisons between pipelines challenging. Diffusion MRI results have been compared to tract tracing in animals (Calabrese et al., 2015; Girard et al., 2020) and to simulations (Farrher et al., 2012; Maier-Hein et al., 2017), but the field is yet to converge on a gold standard pipeline.

Finally, our analysis focused on group connectomes, as these are most commonly studied in the literature. Recent analyses of functional MRI data have suggested that there is considerable individual variability in network architecture that is behaviorally meaningful (Kong et al., 2019; Levakov et al., 2021; Sun et al., 2022). Developing better ways of capturing biologically meaningful individual differences, as distinct from measurement noise, remains an important challenge for the field.

## Conclusions

Our findings indicate that different processing choices affect inferences about network hubs, and that evidence for a concentration of connectivity in hubs occurs in a minor fraction of pipeline variations. Thus, our analysis suggests that it can be quite difficult to identify network hubs in a consistent way, at least across different tractography algorithms and parcellations. However, not all pipeline choices are equal. Although no gold standard pipeline currently exists, some choices are preferred over others. Some denoising procedures, such as the use of eddy correction with outlier replacement and within-slice motion correction, show excellent denoising performance, and are thus recommended (Oldham et al., 2020). ACT (Smith et al., 2012) represents a principled, reasonable constraint on tractography that can be used to remove biologically implausible streamlines. Furthermore, certain parcellations yield parcels that are more functionally homogeneous than others, supporting their biological validity. In this respect, the Schaefer parcellations generally perform quite well with respect to diverse benchmarks (Bryce et al., 2021; Schaefer et al., 2018). However, whether one should choose deterministic or probabilistic tractography is a difficult question to answer definitively. Deterministic tractography is more conservative, but may miss real long-range connections that are important for mapping hub connectivity (Arnatkevičiūtė et al., 2021; Fulcher & Fornito, 2016; van den Heuvel et al., 2012). Probabilistic tractography is better able to resolve such connections but may be prone to false positives. Choices related to different seeding strategies may require more detailed investigation. The incorporation and improvement of sparsity constraints and filtering techniques (Schiavi et al., 2020; Smith et al., 2015b) will be important for improving the accuracy of these approaches. Ongoing assessment with respect to plausible phantoms may help adjudicate between these alternatives (e.g., Maier-Hein et al., 2017). Until then, investigators should assess the robustness of their results by analyzing dMRI data using multiple pipelines, and should be aware of the effects that the choices they exercise in processing their data have on their final results.

## 3. Methods

### 3.1. Participants

294 healthy participants (mean age 23.12 ± 5.18 years, 162 females) were recruited at Monash University with informed consent. All participants self-reported right-handedness and had reported no significant neurological/psychiatric history (i.e., no personal history of neurological or psychiatric disorders, no loss of consciousness or memory due to head injury, and no history of drug use disorder). Further information on sample characteristics is provided elsewhere (Sabaroedin et al., 2019). The study was conducted in accordance with the Monash University Human Research Ethics Committee (reference number 2012001562).

### 3.2. Image Acquisition

T1-weighted (T1w) and diffusion MRIs were acquired on a Siemens (Munich, Germany) Skyra 3T scanner with a 32-channel head coil at Monash Biomedical Imaging in Clayton, Victoria, Australia. T1w structural scans were acquired with the following parameters: 1 mm3 isotropic voxels, TR = 2300 ms, TE = 2.07 ms, TI = 900 ms, and a FOV of 256 mm. Diffusion scans were obtained using an interleaving acquisition with the following parameters: 2.5 mm3 isotropic voxels, TR = 8800 ms, TE = 110 ms, FOV of 240 mm, 60 directions with b = 3000 s/mm2, and seven b = 0 s/mm2 vol. In addition, a single b = 0 s/mm2 was obtained with reversed phase encoding direction for susceptibility field estimation.

### 3.3. Image processing common to all pipelines

Imaging data were processed using the Multi-modal Australian ScienceS Imaging and Visualisation Environment (MASSIVE) high performance computing infrastructure (Goscinski et al., 2014) as described by (Oldham et al., 2020). The analysis evaluated the efficacy of 240 different diffusion MRI processing pipelines in mitigating motion-related artifacts in connectivity estimates, with the pipelines generated by varying choices at each of seven steps (distortion correction, tractography algorithm, propagation constraints, streamline seeding, tractogram re-weighting, edge weighting, and parcellation). We adopted recommendations of Oldham et al. (2020) for three of these (distortion correction, tractogram re-weighting, and edge weighting), as specific options in these steps were shown to reduce the correlation between head movement and structural connectivity. We evaluated effects of the four remaining factors, three of which pertain to the tractography algorithm (probabilistic versus deterministic algorithm, propagation constraints, streamline seeding) and the last of which pertains to parcellation. We further considered how these steps interact with different thresholding and group-aggregation methods. A visual schematic of our pipeline variations is presented in Figure 1. Further details about the choices made at each step are provided in the following sections.

#### 3.3.1. DWI and T1w preprocessing

MRtrix version 3.0.15 (Tournier et al., 2019) and FSL version 5.0.11 (Jenkinson et al., 2012) were used to process the diffusion MRI data. First, FSL’s *topup* was used to estimate the susceptibility-induced off-resonance field using the forward and reverse phase-encoded b = 0 s/mm2 images (Andersson et al., 2003; Smith et al., 2004). Then, FSL’s *eddy* tool was used for motion and eddy current correction, which has been shown to successfully mitigate motion-related artifact in connectivity estimates (Oldham et al., 2020), and which incorporates both (i) a Gaussian process-based generative model for volume prediction and realignment (Andersson & Sotiropoulos, 2016) and (ii) reconstruction and replacement of slices with significant signal dropout (Andersson et al., 2016, 2017). The following parameters were used for slice-to-volume correction: temporal order of movement = 30, iterations = 5, strength of temporal regularization = 6, and trilinear interpolation. Finally, FAST in FSL was used to correct for B1 field inhomogeneities (Smith et al., 2004; Zhang et al., 2001).

The diffusion images were then co-registered to the T1w images via a rigid-body transformation using FSL’s *FLIRT* (Jenkinson et al., 2002; Jenkinson & Smith, 2001) and the inverse of this transformation was used to map the T1w image to the subject’s native diffusion space, where all tractography was performed. FreeSurfer version 5.3 (Fischl, 2012) was used to extract cortical surface models (grey/white matter surface and grey/CSF surface) from T1w images. All outputs were visually inspected and manually corrected, if required. Parcellation schemes (detailed in 3.4.4) were applied to the cortical surface models; these were then projected to the T1w image grid and used to define network nodes.

### 3.4. Pipeline-specific image processing

In this section, we outline the key pipeline variations considered in our analysis.

#### 3.4.1. Streamline seeding algorithm

Streamline seeding is the process by which voxels are selected to be the propagation points for streamlines. As in Oldham et al., (2020), we compare three streamline seeding algorithms:

1. White matter (WM): voxels coded as white matter are randomly chosen as streamline seeds.
2. Grey matter-white matter interface (GMWMI): voxels containing a gradient between grey matter and white matter are chosen as streamline seeds, with the aim of improving the tractography of shorter fibers (Smith et al., 2013, 2015a).
3. Dynamic: the relative difference between the predicted fiber density (based on the diffusion model) and the current density is used to inform the probability of choosing a particular location as a seed, with the aim of correcting for under- or over-sampling of a given fiber tract (Smith et al., 2015b).

#### 3.4.2. Streamline tractography algorithm

Most tractography algorithms are classified as being either deterministic or probabilistic. Deterministic algorithms tend to be more conservative and thus prone to false negatives, whilst probabilistic algorithms are more sensitive but can be prone to false positives (Reveley et al., 2015; Sarwar et al., 2019; Thomas et al., 2014). We compared an exemplar of each class, both of which were implemented in MRtrix3 (Tournier et al., 2019):

1. Deterministic tractography was performed using the Fibre Assignment by Continuous Tractography (FACT) algorithm (Mori et al., 1999; Mori & van Zijl, 2002).
2. Probabilistic tractography was performed using Second-order integration over Fibre Orientation Distributions (iFOD2) (Tournier et al., 2007, 2010, 2012).

For both tractography algorithms, 2,000,000 streamlines were generated with a maximum length of 400 mm, a maximum curvature of 45° per step, the default step size (1.25 mm for FACT; 0.25 mm for iFOD2), and the default termination criterion (0.05 amplitude of the primary eigenvector for FACT; 0.05 FOD amplitude for iFOD2).

#### 3.4.3. Streamline propagation constraint

Tractography algorithms often track streamlines through anatomically implausible areas (e.g., ventricles), which can be addressed by imposing some constraints on streamline propagation. We examined two spatial constraints:

1. Grey and white matter masking (GWM), involving the use of a binary mask (combining the grey and white matter masks from the FreeSurfer segmentation) that ensures streamlines only travel through brain parenchyma.
2. Anatomically Constrained Tractography (ACT), which uses a multi-tissue segmentation (cortical grey matter, subcortical grey matter, white matter, and CSF) and a series of propagation rules to ensure that streamlines follow anatomically viable paths (Smith et al., 2012).

Because the implementation of GMWMI seeding in MRtrix3 requires ACT, pipelines combining GWM and GMWMI were excluded. The above combinations therefore resulted in a total of ten different tractography workflows for comparison.

#### 3.4.4. Parcellation

A wide variety of parcellations have been used in the connectomics literature (Arslan et al., 2018; de Reus & van den Heuvel, 2013b; Lawrence et al., 2021), which can affect various network properties (Eickhoff et al., 2015; Fornito et al., 2010; Zalesky et al., 2010). We compared four different cortical parcellation schemes derived using three different approaches:

1. The Desikan-Killiany parcellation (DK68), comprising 34 cortical nodes in each hemisphere delineated using sulcal and gyral landmarks (Desikan et al., 2006).
2. The Human Connectome Project MMP1 parcellation (HCP360), comprising 180 cortical nodes per hemisphere defined using a semi-automated pipeline that leverages information on regional cortical architecture, function, connectivity, and topography (Glasser et al., 2016).
3. The Schaefer et al. (Schaefer et al., 2018) 200 and 500 node parcellations (S200 and S500), generated based on local gradients of global profile similarities in regional functional coupling estimates.

These parcellations represent both (i) distinct technical and methodological approaches relying different biological properties; and (ii) diversity in the sizes and shapes of parcels produced. Each parcellation was generated using surface models estimated by FreeSurfer using *fsaverage* coordinates; these were registered to each individual’s surface and then projected out to the T1w volume. The combination of ten tractography workflows and four parcellations resulted in a total of 40 pipelines for reconstructing individual connectomes.

#### 3.4.5. Group aggregation

Having generated individual connectomes using the above parameters, we compared four methods for aggregating the data to obtain a group-representative connectome:

1. Edge weight, which retains edges with the largest mean weight, up to a specified density.
2. Edge coefficient of variation (CV), which retains edges with the smallest CV across participants (Roberts et al., 2017), up to a specified density.
3. Edge consistency, which retains edges that are present (i.e., with non-zero weight) in the greatest number of participants (de Reus & van den Heuvel, 2013a), up to a specified density. Whilst this can be formulated by selecting consistency thresholds which must be met, here we equivalently specify density thresholds and retain the most consistent edges to ensure that connectomes are density-matched, which facilitates comparisons across pipelines (see 3.4.6).
4. Edge distance-dependent binning, which bins edges according to their length, using a specified number of bins, and retains edges that are most frequently present within each bin (Betzel et al., 2019).

Note that for each method, the final weight of the retained edges is equal to the mean of the edge weights across all participants; it is only the choice of which edges to retain that changes.

#### 3.4.6. Group thresholding

Having generated a group connectome using one of the above approaches, we thresholded the resulting matrix at different levels using one of two approaches, depending on the aggregation method:

1. Density thresholds were used for group connectomes aggregated using edge weight, edge CV, and edge consistency, retaining the top-ranked edges according to each measure, evaluating densities spanning 5% to 30%, in increments of 2.5%.
2. The number of bins was used for the group connectome generated with edge distance-dependent binning, in which we changed the number of bins from 10 to 110, in increments of 10. In general, increasing the number of bins increases the density of the group connectome, resulting in networks with densities spanning 2% to 94%. Note that connectomes generated in this way were evaluated separately when evaluating how network properties depend on connectome density.

The combination of four group aggregation methods and eleven thresholds for each threshold resulted in a total of 44 group reconstruction regimes for comparison.

### 3.5. Statistical analysis

We first evaluated how the above processing choices affect properties of the degree distribution of the connectome. The degree distribution defines the extent to which connectivity is concentrated in network hubs. Distributions with a heavy tail imply the existence of highly connected hubs, whereas distributions with an approximately exponential fall-off imply that the concentration of connections on putative hubs does not exceed the expectations of a random network (Fornito et al., 2016). We therefore concentrated on the properties of the distribution tails. Distributions of both binary and weighted node degree in brain networks have been previously described as heavy-tailed (taken here to mean that the tail decays sub-exponentially), but the precise distribution they follow has been the subject of debate (Buzsáki & Mizuseki, 2014; Fornito et al., 2016; Roberts et al., 2015; Zucca et al., 2019). Moreover, parametric modelling of these distributions is dependent on user-defined inputs, such as the choice of the models under consideration or the model fitting procedures used, resulting in another source of variability when comparing computational pipelines.

Fitting the empirical degree distribution to the generalized extreme value distribution and obtaining a tail-decay index can mitigate these problems (Gomes & Guillou, 2015; Haan & Ferreira, 2006; Hill, 1975). However, this approach often requires a large number of data points (Németh & Zempléni, 2020) and still depends on heuristic measures to define the start and end of the tail (Bauke, 2007; Gomes et al., 2009; Paulauskas & Vaičiulis, 2017). We therefore used the non-parametric approach described by (Jordanova & Petkova, 2017), which more directly focuses on the question of heavy-tailedness.

In brief, to determine if the distribution of a random variable *X* has a heavier right tail than the exponential distribution, we calculate the empirical first and third quartiles, 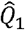 and 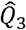 respectively, and the interquartile range, 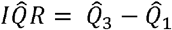 We then define the “right-tailedness” of the distribution as the probability that a random drawn observation from the distribution is greater than the value given by 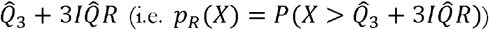, utilizing the commonly used definition of extreme outliers (McGill et al., 1978). This value can be compared to the right-tailedness of the exponential distribution (*X* ∼ e^-*λX*^), which is invariant to the shape parameter *λ*, such that *pR* (*X*) exp(-*λ* • ln(3^3^ • 4)/ *λ*) 1/108 ≈ 0.009 (Jordanova & Petkova, 2017). This analytic solution offers a convenient threshold for determining whether a distribution has a heavier right tail than the exponential distribution, with heavy-tailedness implied if the empirical *p*_*R*_ > 0.009.

Additionally, we quantified the asymmetry of the whole distribution using the skewness (the third standardized moment), which is also constant for the exponential distribution (skewness = 2). Finally, for completeness, we calculated the excess kurtosis (the fourth standardized moment), which provides an alternative method for capturing the behavior at the tails (DeCarlo, 1997; Westfall, 2014). This measure has been shown to be robust for detecting outliers in small samples (Hayes et al., 2007; Livesey, 2007) and is also independent of the shape parameter of the exponential distribution (excess kurtosis = 6). We note that other methods are available, including tail index estimation (Caers & Dyck, 1998; Németh & Zempléni, 2020), parametric fitting (Zucca et al., 2019), and skewness-free kurtosis measures (Critchley & Jones, 2008; Eberl & Klar, 2022; Jones et al., 2011; Oja, 1981). However, as with parametric modelling, these methods rely on user-defined algorithms or parameters (such as the number of quantiles to be used or the cut-off point for initialization of the tail), similarly making comparisons difficult.

After characterizing the statistical properties of the degree distribution, we examined the spatial distribution of inter-regional connectivity by considering the degree sequence. The degree sequence encodes the assignment of degree values to specific nodes, hence capturing the spatial position or topography of network hubs. Within parcellations, we compared the consistency of hub topography and the effects of surface area on hubness. Between parcellations, we examined the consistency of hub topography across different pipelines qualitatively, as the lack of region-to-region correspondence precludes a direct comparison.

## Acknowledgements

AF was supported by the Sylvia and Charles Viertel Charitable Foundation, the National Health and Medical Research Council (IDs: 1149292 and 1197431), and the Australian Research Council (IDs: DP200103509). MAB is supported by a NHMRC Senior Research Fellowship (Level B).

## CRediT authorship contribution statement

**Mehul Gajwani:** Conceptualization, Methodology, Formal analysis, Writing – original draft. **Stuart Oldham:** Conceptualization, Formal analysis, Methodology, Software, Writing – review & editing. **James C. Pang:** Supervision, Writing – review & editing. **Aurina Arnatkevic**□**iūtė:** Methodology, Writing – review & editing. **Jeggan Tiego:** Resources, Writing – review & editing. **Mark A. Bellgrove:** Funding acquisition, Writing – review & editing. **Alex Fornito:** Conceptualization, Methodology, Formal analysis, Writing – original draft, Supervision, Funding acquisition.

## References

Andersson, J. L. R., Graham, M. S., Drobnjak, I., Zhang, H., Filippini, N., & Bastiani, M. (2017). Towards a comprehensive framework for movement and distortion correction of diffusion MR images: Within volume movement. NeuroImage, 152, 450–466. https://doi.org/10.1016/j.neuroimage.2017.02.085

Andersson, J. L. R., Graham, M. S., Zsoldos, E., & Sotiropoulos, S. N. (2016). Incorporating outlier detection and replacement into a non-parametric framework for movement and distortion correction of diffusion MR images. NeuroImage, 141, 556–572. https://doi.org/10.1016/j.neuroimage.2016.06.058

Andersson, J. L. R., Skare, S., & Ashburner, J. (2003). How to correct susceptibility distortions in spin-echo echo-planar images: Application to diffusion tensor imaging. NeuroImage, 20(2), 870–888. https://doi.org/10.1016/S1053-8119(03)00336-7

Andersson, J. L. R., & Sotiropoulos, S. N. (2016). An integrated approach to correction for off-resonance effects and subject movement in diffusion MR imaging. NeuroImage, 125, 1063–1078. https://doi.org/10.1016/j.neuroimage.2015.10.019

Arnatkevičiūtė, A., Fulcher, B. D., & Fornito, A. (2019). Uncovering the Transcriptional Correlates of Hub Connectivity in Neural Networks. Frontiers in Neural Circuits, 13, 47. https://doi.org/10.3389/fncir.2019.00047

Arnatkevičiūtė, A., Fulcher, B. D., Oldham, S., Tiego, J., Paquola, C., Gerring, Z., Aquino, K., Hawi, Z., Johnson, B., Ball, G., Klein, M., Deco, G., Franke, B., Bellgrove, M. A., & Fornito, A. (2021). Genetic influences on hub connectivity of the human connectome. Nature Communications, 12(1), 4237. https://doi.org/10.1038/s41467-021-24306-2

Arnatkevic□iūtė, A., Fulcher, B. D., Pocock, R., & Fornito, A. (2018). Hub connectivity, neuronal diversity, and gene expression in the Caenorhabditis elegans connectome. PLoS Computational Biology, 14(2), e1005989. https://doi.org/10.1371/journal.pcbi.1005989

Arslan, S., Ktena, S. I., Makropoulos, A., Robinson, E. C., Rueckert, D., & Parisot, S. (2018). Human brain mapping: A systematic comparison of parcellation methods for the human cerebral cortex. NeuroImage, 170, 5–30. https://doi.org/10.1016/j.neuroimage.2017.04.014

Bauke, H. (2007). Parameter estimation for power-law distributions by maximum likelihood methods. The European Physical Journal B, 58(2), 167–173. https://doi.org/10.1140/epjb/e2007-00219-y

Betzel, R. F., & Bassett, D. S. (2017). Multi-scale brain networks. NeuroImage, 160, 73–83. https://doi.org/10.1016/j.neuroimage.2016.11.006

Betzel, R. F., Griffa, A., Hagmann, P., & Mišić, B. (2019). Distance-dependent consensus thresholds for generating group-representative structural brain networks. Network Neuroscience, 3(2), 475–496. https://doi.org/10.1162/netn_a_00075

Bordier, C., Nicolini, C., & Bifone, A. (2017). Graph Analysis and Modularity of Brain Functional Connectivity Networks: Searching for the Optimal Threshold. Frontiers in Neuroscience, 11. https://www.frontiersin.org/articles/10.3389/fnins.2017.00441

Brodmann, K. (n.d.). Vergleichende Lokalisationslehre der Grosshirnrinde in ihren Prinzipien dargestellt auf Grund des Zellenbaues / [K. Brodmann]. Retrieved 5 October 2022, from https://jstor.org/stable/community.24783456

Bryce, N. V., Flournoy, J. C., Guassi Moreira, J. F., Rosen, M. L., Sambook, K. A., Mair, P., & McLaughlin, K. A. (2021). Brain parcellation selection: An overlooked decision point with meaningful effects on individual differences in resting-state functional connectivity. NeuroImage, 243, 118487. https://doi.org/10.1016/j.neuroimage.2021.118487

Buzsáki, G., & Mizuseki, K. (2014). The log-dynamic brain: How skewed distributions affect network operations. Nature Reviews Neuroscience, 15(4), Article 4. https://doi.org/10.1038/nrn3687

Caers, J., & Dyck, J. V. (1998). Nonparametric tail estimation using a double bootstrap method. Computational Statistics & Data Analysis, 29(2), 191– 211. https://doi.org/10.1016/S0167-9473(98)00060-7

Calabrese, E., Badea, A., Cofer, G., Qi, Y., & Johnson, G. A. (2015). A Diffusion MRI Tractography Connectome of the Mouse Brain and Comparison with Neuronal Tracer Data. Cerebral Cortex (New York, N.Y.: 1991), 25(11), 4628–4637. https://doi.org/10.1093/cercor/bhv121

Clauset, A., Shalizi, C. R., & Newman, M. E. J. (2009). Power-Law Distributions in Empirical Data. SIAM Review, 51(4), 661–703. https://doi.org/10.1137/070710111

Critchley, F., & Jones, M. C. (2008). Asymmetry and Gradient Asymmetry Functions: Density-Based Skewness and Kurtosis. Scandinavian Journal of Statistics, 35(3), 415–437. https://doi.org/10.1111/j.1467-9469.2008.00599.x

Crossley, N. A., Mechelli, A., Scott, J., Carletti, F., Fox, P. T., McGuire, P., & Bullmore, E. T. (2014). The hubs of the human connectome are generally implicated in the anatomy of brain disorders. Brain: A Journal of Neurology, 137(Pt 8), 2382–2395. https://doi.org/10.1093/brain/awu132

de Lange, S. C., Scholtens, L. H., van den Berg, L. H., Boks, M. P., Bozzali, M., Cahn, W., Dannlowski, U., Durston, S., Geuze, E., van Haren, N. E. M., Hillegers, M. H. J., Koch, K., Jurado, M. Á., Mancini, M., Marqués-Iturria, I., Meinert, S., Ophoff, R. A., Reess, T. J., Repple, J., … van den Heuvel, M. P. (2019). Shared vulnerability for connectome alterations across psychiatric and neurological brain disorders. Nature Human Behaviour, 3(9), Article 9. https://doi.org/10.1038/s41562-019-0659-6

de Reus, M. A., & van den Heuvel, M. P. (2013a). Estimating false positives and negatives in brain networks. NeuroImage, 70, 402–409. https://doi.org/10.1016/j.neuroimage.2012.12.066

de Reus, M. A., & van den Heuvel, M. P. (2013b). The parcellation-based connectome: Limitations and extensions. NeuroImage, 80, 397–404. https://doi.org/10.1016/j.neuroimage.2013.03.053

DeCarlo, L. T. (1997). On the Meaning and Use of Kurtosis. Psychological Methods, 2, 292–307.

Desikan, R. S., Ségonne, F., Fischl, B., Quinn, B. T., Dickerson, B. C., Blacker, D., Buckner, R. L., Dale, A. M., Maguire, R. P., Hyman, B. T., Albert, M. S., & Killiany, R. J. (2006). An automated labeling system for subdividing the human cerebral cortex on MRI scans into gyral based regions of interest. NeuroImage, 31(3), 968–980. https://doi.org/10.1016/j.neuroimage.2006.01.021

Eberl, A., & Klar, B. (2022). Centre-free kurtosis orderings for asymmetric distributions (arXiv:2210.04850). arXiv. http://arxiv.org/abs/2210.04850

Eickhoff, S. B., Thirion, B., Varoquaux, G., & Bzdok, D. (2015). Connectivity-based parcellation: Critique and implications. Human Brain Mapping, 36(12), 4771–4792. https://doi.org/10.1002/hbm.22933

Fagerholm, E. D., Hellyer, P. J., Scott, G., Leech, R., & Sharp, D. J. (2015). Disconnection of network hubs and cognitive impairment after traumatic brain injury. Brain, 138(6), 1696–1709. https://doi.org/10.1093/brain/awv075

Fan, Y., Shi, F., Smith, J. K., Lin, W., Gilmore, J. H., & Shen, D. (2011). Brain anatomical networks in early human brain development. NeuroImage, 54(3), 1862–1871. https://doi.org/10.1016/j.neuroimage.2010.07.025

Farrher, E., Kaffanke, J., Celik, A. A., Stöcker, T., Grinberg, F., & Shah, N. J. (2012). Novel multisection design of anisotropic diffusion phantoms. Magnetic Resonance Imaging, 30(4), 518–526. https://doi.org/10.1016/j.mri.2011.12.012

Fischl, B. (2012). FreeSurfer. NeuroImage, 62(2), 774–781. https://doi.org/10.1016/j.neuroimage.2012.01.021

Fornito, A., Zalesky, A., & Breakspear, M. (2015). The connectomics of brain disorders. Nature Reviews Neuroscience, 16(3), Article 3. https://doi.org/10.1038/nrn3901

Fornito, A., Zalesky, A., & Bullmore, E. (2016). Fundamentals of Brain Network Analysis (1st ed.). Academic Press. https://doi.org/10.1016/B978-0-12-407908-3.00001-7

Fornito, A., Zalesky, A., & Bullmore, E. T. (2010). Network scaling effects in graph analytic studies of human resting-state FMRI data. Frontiers in Systems Neuroscience, 4, 22. https://doi.org/10.3389/fnsys.2010.00022

Fulcher, B. D., & Fornito, A. (2016). A transcriptional signature of hub connectivity in the mouse connectome. Proceedings of the National Academy of Sciences of the United States of America, 113(5), 1435–1440. https://doi.org/10.1073/pnas.1513302113

Genon, S., Reid, A., Li, H., Fan, L., Müller, V. I., Cieslik, E. C., Hoffstaedter, F., Langner, R., Grefkes, C., Laird, A. R., Fox, P. T., Jiang, T., Amunts, K., & Eickhoff, S. B. (2018). The heterogeneity of the left dorsal premotor cortex evidenced by multimodal connectivity-based parcellation and functional characterization. NeuroImage, 170, 400–411. https://doi.org/10.1016/j.neuroimage.2017.02.034

Girard, G., Caminiti, R., Battaglia-Mayer, A., St-Onge, E., Ambrosen, K. S., Eskildsen, S. F., Krug, K., Dyrby, T. B., Descoteaux, M., Thiran, J.-P., & Innocenti, G. M. (2020). On the cortical connectivity in the macaque brain: A comparison of diffusion tractography and histological tracing data. NeuroImage, 221, 117201. https://doi.org/10.1016/j.neuroimage.2020.117201

Glasser, M. F., Coalson, T. S., Robinson, E. C., Hacker, C. D., Harwell, J., Yacoub, E., Ugurbil, K., Andersson, J., Beckmann, C. F., Jenkinson, M., Smith, S. M., & Van Essen, D. C. (2016). A multi-modal parcellation of human cerebral cortex. Nature, 536(7615), Article 7615. https://doi.org/10.1038/nature18933

Gollo, L. L., Roberts, J. A., Cropley, V. L., Di Biase, M. A., Pantelis, C., Zalesky, A., & Breakspear, M. (2018). Fragility and volatility of structural hubs in the human connectome. Nature Neuroscience, 21(8), Article 8. https://doi.org/10.1038/s41593-018-0188-z

Gomes, M. I., & Guillou, A. (2015). Extreme Value Theory and Statistics of Univariate Extremes: A Review. International Statistical Review, 83(2), 263– 292. https://doi.org/10.1111/insr.12058

Gomes, M. I., Pestana, D., & Caeiro, F. (2009). A note on the asymptotic variance at optimal levels of a bias-corrected Hill estimator. Statistics & Probability Letters, 79(3), 295–303. https://doi.org/10.1016/j.spl.2008.08.016

Gonzalez-Aquines, A., Moreno-Andrade, T., Gongora-Rivera, F., Cordero-Perez, A. C., Ortiz-Jiménez, X., Cavazos-Luna, O., Garza-Villareal, E., Campos-Coy, M., & Elizondo-Riojas, G. (2019). The Role of Tractography in Ischemic Stroke: A Review of the Literature. Revista Medicina Universitaria, 20(4), 1597. https://doi.org/10.24875/RMU.18000021

Goscinski, W. J., McIntosh, P., Felzmann, U. C., Maksimenko, A., Hall, C. J., Gureyev, T., Thompson, D., Janke, A., Galloway, G., Killeen, N. E. B., Raniga, P., Kaluza, O., Ng, A., Poudel, G., Barnes, D., Nguyen, T., Bonnington, P., & Egan, G. F. (2014). The multi-modal Australian ScienceS Imaging and Visualization Environment (MASSIVE) high performance computing infrastructure: Applications in neuroscience and neuroinformatics research. Frontiers in Neuroinformatics, 0. https://doi.org/10.3389/fninf.2014.00030

Haan, L. de, & Ferreira, A. (2006). Extreme value theory: An introduction. Springer.

Hayes, K., Kinsella, A., & Coffey, N. (2007). A note on the use of outlier criteria in Ontario laboratory quality control schemes. Clinical Biochemistry, 40(3), 147–152. https://doi.org/10.1016/j.clinbiochem.2006.08.019

Hill, B. M. (1975). A Simple General Approach to Inference About the Tail of a Distribution. The Annals of Statistics, 3(5), 1163–1174. https://doi.org/10.1214/aos/1176343247

Horbruegger, M., Loewe, K., Kaufmann, J., Wagner, M., Schippling, S., Pawlitzki, M., & Schoenfeld, M. A. (2019). Anatomically constrained tractography facilitates biologically plausible fiber reconstruction of the optic radiation in multiple sclerosis. NeuroImageLI: Clinical, 22, 101740. https://doi.org/10.1016/j.nicl.2019.101740

Jenkinson, M., Bannister, P., Brady, M., & Smith, S. (2002). Improved optimization for the robust and accurate linear registration and motion correction of brain images. NeuroImage, 17(2), 825–841. https://doi.org/10.1016/s1053-8119(02)91132-8

Jenkinson, M., Beckmann, C. F., Behrens, T. E. J., Woolrich, M. W., & Smith, S. M. (2012). FSL. NeuroImage, 62(2), 782–790. https://doi.org/10.1016/j.neuroimage.2011.09.015

Jenkinson, M., & Smith, S. (2001). A global optimisation method for robust affine registration of brain images. Medical Image Analysis, 5(2), 143–156. https://doi.org/10.1016/s1361-8415(01)00036-6

Jeurissen, B., Descoteaux, M., Mori, S., & Leemans, A. (2019). Diffusion MRI fiber tractography of the brain. NMR in Biomedicine, 32(4), e3785. https://doi.org/10.1002/nbm.3785

Jones, D. K., & Cercignani, M. (2010). Twenty-five pitfalls in the analysis of diffusion MRI data. NMR in Biomedicine, 23(7), 803–820. https://doi.org/10.1002/nbm.1543

Jones, M. C., Rosco, J. F., & Pewsey, A. (2011). Skewness-Invariant Measures of Kurtosis. The American Statistician, 65(2), 89–95. https://doi.org/10.1198/tast.2011.10194

Jordanova, P. K., & Petkova, M. P. (2017). Measuring heavy-tailedness of distributions. AIP Conference Proceedings, 1910(1), 060002. https://doi.org/10.1063/1.5013996

Kong, R., Li, J., Orban, C., Sabuncu, M. R., Liu, H., Schaefer, A., Sun, N., Zuo, X.-N., Holmes, A. J., Eickhoff, S. B., & Yeo, B. T. T. (2019). Spatial Topography of Individual-Specific Cortical Networks Predicts Human Cognition, Personality, and Emotion. Cerebral Cortex, 29(6), 2533– 2551. https://doi.org/10.1093/cercor/bhy123

Lawrence, R. M., Bridgeford, E. W., Myers, P. E., Arvapalli, G. C., Ramachandran, S. C., Pisner, D. A., Frank, P. F., Lemmer, A. D., Nikolaidis, A., & Vogelstein, J. T. (2021). Standardizing human brain parcellations. Scientific Data, 8(1), Article 1. https://doi.org/10.1038/s41597-021-00849-3

Levakov, G., Faskowitz, J., Avidan, G., & Sporns, O. (2021). Mapping individual differences across brain network structure to function and behavior with connectome embedding. NeuroImage, 242, 118469. https://doi.org/10.1016/j.neuroimage.2021.118469

Li, L., Rilling, J. K., Preuss, T. M., Glasser, M. F., & Hu, X. (2012). The effects of connection reconstruction method on the interregional connectivity of brain networks via diffusion tractography. Human Brain Mapping, 33(8), 1894–1913. https://doi.org/10.1002/hbm.21332

Lilja, Y., Ljungberg, M., Starck, G., Malmgren, K., Rydenhag, B., & Nilsson, D. T. (2014). Visualizing Meyer’s loop: A comparison of deterministic and probabilistic tractography. Epilepsy Research, 108(3), 481–490. https://doi.org/10.1016/j.eplepsyres.2014.01.017

Lipp, I., Parker, G. D., Tallantyre, E. C., Goodall, A., Grama, S., Patitucci, E., Heveron, P., Tomassini, V., & Jones, D. K. (2020). Tractography in the presence of multiple sclerosis lesions. NeuroImage, 209, 116471. https://doi.org/10.1016/j.neuroimage.2019.116471

Livesey, J. H. (2007). Kurtosis provides a good omnibus test for outliers in small samples. Clinical Biochemistry, 40(13), 1032–1036. https://doi.org/10.1016/j.clinbiochem.2007.04.003

Maier-Hein, K. H., Neher, P. F., Houde, J.-C., Côté, M.-A., Garyfallidis, E., Zhong, J., Chamberland, M., Yeh, F.-C., Lin, Y.-C., Ji, Q., Reddick, W. E., Glass, J. O., Chen, D. Q., Feng, Y., Gao, C., Wu, Y., Ma, J., He, R., Li, Q., … Descoteaux, M. (2017). The challenge of mapping the human connectome based on diffusion tractography. Nature Communications, 8(1), Article 1. https://doi.org/10.1038/s41467-017-01285-x

Martínez-Heras, E., Varriano, F., Prčkovska, V., Laredo, C., Andorrà, M., Martínez-Lapiscina, E. H., Calvo, A., Lampert, E., Villoslada, P., Saiz, A., Prats-Galino, A., & Llufriu, S. (2015). Improved Framework for Tractography Reconstruction of the Optic Radiation. PLOS ONE, 10(9), e0137064. https://doi.org/10.1371/journal.pone.0137064

McGill, R., Tukey, J. W., & Larsen, W. A. (1978). Variations of Box Plots. The American Statistician, 32(1), 12–16. https://doi.org/10.2307/2683468

Mišić, B., Betzel, R. F., Nematzadeh, A., Goñi, J., Griffa, A., Hagmann, P., Flammini, A., Ahn, Y.-Y., & Sporns, O. (2015). Cooperative and Competitive Spreading Dynamics on the Human Connectome. Neuron, 86(6), 1518–1529. https://doi.org/10.1016/j.neuron.2015.05.035

Mochizuki, M., Uchiyama, Y., Domen, K., & Koyama, T. (2023). Applicability of automated tractography during acute care stroke rehabilitation. Journal of Physical Therapy Science, 35(2), 156–162. https://doi.org/10.1589/jpts.35.156

Modha, D. S., & Singh, R. (2010). Network architecture of the long-distance pathways in the macaque brain. Proceedings of the National Academy of Sciences, 107(30), 13485–13490. https://doi.org/10.1073/pnas.1008054107

Mori, S., Crain, B. J., Chacko, V. P., & Van Zijl, P. C. M. (1999). Three-dimensional tracking of axonal projections in the brain by magnetic resonance imaging. Annals of Neurology, 45(2), 265–269. https://doi.org/10.1002/1531-8249(199902)45:2<265::AID-ANA21>3.0.CO;2-3

Mori, S., & van Zijl, P. C. M. (2002). Fiber tracking: Principles and strategies – a technical review. NMR in Biomedicine, 15(7–8), 468–480. https://doi.org/10.1002/nbm.781

Németh, L., & Zempléni, A. (2020). Regression Estimator for the Tail Index. Journal of Statistical Theory and Practice, 14(3), 48. https://doi.org/10.1007/s42519-020-00114-7

Oja, H. (1981). On Location, Scale, Skewness and Kurtosis of Univariate Distributions. Scandinavian Journal of Statistics, 8(3), 154–168.

Oldham, S., Arnatkevic□iūtė, A., Smith, R. E., Tiego, J., Bellgrove, M. A., & Fornito, A. (2020). The efficacy of different preprocessing steps in reducing motion-related confounds in diffusion MRI connectomics. NeuroImage, 222, 117252. https://doi.org/10.1016/j.neuroimage.2020.117252

Oldham, S., & Fornito, A. (2019). The development of brain network hubs. Developmental Cognitive Neuroscience, 36, 100607. https://doi.org/10.1016/j.dcn.2018.12.005

Oldham, S., Fulcher, B. D., Aquino, K., Arnatkevičiūtė, A., Paquola, C., Shishegar, R., & Fornito, A. (2022). Modeling spatial, developmental, physiological, and topological constraints on human brain connectivity. Science Advances, 8(22), eabm6127. https://doi.org/10.1126/sciadv.abm6127

Oldham, S., Fulcher, B., Parkes, L., Arnatkevic Iūtė, A., Suo, C., & Fornito, A. (2019). Consistency and differences between centrality measures across distinct classes of networks. PloS One, 14(7), e0220061. https://doi.org/10.1371/journal.pone.0220061

Paulauskas, V., & Vaičiulis, M. (2017). A class of new tail index estimators. Annals of the Institute of Statistical Mathematics, 69(2), 461–487. https://doi.org/10.1007/s10463-015-0548-3

Reveley, C., Seth, A. K., Pierpaoli, C., Silva, A. C., Yu, D., Saunders, R. C., Leopold, D. A., & Ye, F. Q. (2015). Superficial white matter fiber systems impede detection of long-range cortical connections in diffusion MR tractography. Proceedings of the National Academy of Sciences, 112(21), E2820–E2828. https://doi.org/10.1073/pnas.1418198112

Richards, T. J., Anderson, K. L., & Anderson, J. S. (2021). “Fully automated segmentation of the corticospinal tract using the TractSeg algorithm in patients with brain tumors”. Clinical Neurology and Neurosurgery, 210, 107001. https://doi.org/10.1016/j.clineuro.2021.107001

Roberts, J. A., Boonstra, T. W., & Breakspear, M. (2015). The heavy tail of the human brain. Current Opinion in Neurobiology, 31, 164–172. https://doi.org/10.1016/j.conb.2014.10.014

Roberts, J. A., Perry, A., Roberts, G., Mitchell, P. B., & Breakspear, M. (2017). Consistency-based thresholding of the human connectome. NeuroImage, 145, 118–129. https://doi.org/10.1016/j.neuroimage.2016.09.053

Sabaroedin, K., Tiego, J., Parkes, L., Sforazzini, F., Finlay, A., Johnson, B., Pinar, A., Cropley, V., Harrison, B. J., Zalesky, A., Pantelis, C., Bellgrove, M., & Fornito, A. (2019). Functional Connectivity of Corticostriatal Circuitry and Psychosis-like Experiences in the General Community. Biological Psychiatry, 86(1), 16–24. https://doi.org/10.1016/j.biopsych.2019.02.013

Sanvito, F., Caverzasi, E., Riva, M., Jordan, K. M., Blasi, V., Scifo, P., Iadanza, A., Crespi, S. A., Cirillo, S., Casarotti, A., Leonetti, A., Puglisi, G., Grimaldi, M., Bello, L., Gorno-Tempini, M. L., Henry, R. G., Falini, A., & Castellano, A. (2020). FMRI-Targeted High-Angular Resolution Diffusion MR Tractography to Identify Functional Language Tracts in Healthy Controls and Glioma Patients. Frontiers in Neuroscience, 14. https://www.frontiersin.org/articles/10.3389/fnins.2020.00225

Sarwar, T., Ramamohanarao, K., & Zalesky, A. (2019). Mapping connectomes with diffusion MRI: Deterministic or probabilistic tractography? Magnetic Resonance in Medicine, 81(2), 1368–1384. https://doi.org/10.1002/mrm.27471

Sarwar, T., Ramamohanarao, K., & Zalesky, A. (2021). A critical review of connectome validation studies. NMR in Biomedicine, 34(12), e4605. https://doi.org/10.1002/nbm.4605

Schaefer, A., Kong, R., Gordon, E. M., Laumann, T. O., Zuo, X.-N., Holmes, A. J., Eickhoff, S. B., & Yeo, B. T. T. (2018). Local-Global Parcellation of the Human Cerebral Cortex from Intrinsic Functional Connectivity MRI. Cerebral Cortex, 28(9), 3095–3114. https://doi.org/10.1093/cercor/bhx179

Schiavi, S., Ocampo-Pineda, M., Barakovic, M., Petit, L., Descoteaux, M., Thiran, J.-P., & Daducci, A. (2020). A new method for accurate in vivo mapping of human brain connections using microstructural and anatomical information. Science Advances, 6(31), eaba8245. https://doi.org/10.1126/sciadv.aba8245

Schilling, K. G., Petit, L., Rheault, F., Remedios, S., Pierpaoli, C., Anderson, A. W., Landman, B. A., & Descoteaux, M. (2020). Brain connections derived from diffusion MRI tractography can be highly anatomically accurate—If we know where white matter pathways start, where they end, and where they do not go. Brain Structure and Function, 225(8), 2387–2402. https://doi.org/10.1007/s00429-020-02129-z

Shu, N., Liu, Y., Li, K., Duan, Y., Wang, J., Yu, C., Dong, H., Ye, J., & He, Y. (2011). Diffusion Tensor Tractography Reveals Disrupted Topological Efficiency in White Matter Structural Networks in Multiple Sclerosis. Cerebral Cortex, 21(11), 2565–2577. https://doi.org/10.1093/cercor/bhr039

Sleurs, C., Jacobs, S., Counsell, S. J., Christiaens, D., Tournier, J.-D., Sunaert, S., Van Beek, K., Uyttebroeck, A., Deprez, S., Batalle, D., & Lemiere, J. (2021). Brain network hubs and cognitive performance of survivors of childhood infratentorial tumors. Radiotherapy and Oncology, 161, 118– 125. https://doi.org/10.1016/j.radonc.2021.05.028

Smith, R. E., Tournier, J.-D., Calamante, F., & Connelly, A. (2012). Anatomically-constrained tractography: Improved diffusion MRI streamlines tractography through effective use of anatomical information. NeuroImage, 62(3), 1924–1938. https://doi.org/10.1016/j.neuroimage.2012.06.005

Smith, R. E., Tournier, J.-D., Calamante, F., & Connelly, A. (2013). SIFT: Spherical-deconvolution informed filtering of tractograms. NeuroImage, 67, 298–312. https://doi.org/10.1016/j.neuroimage.2012.11.049

Smith, R. E., Tournier, J.-D., Calamante, F., & Connelly, A. (2015a). The effects of SIFT on the reproducibility and biological accuracy of the structural connectome. NeuroImage, 104, 253–265. https://doi.org/10.1016/j.neuroimage.2014.10.004

Smith, R. E., Tournier, J.-D., Calamante, F., & Connelly, A. (2015b). SIFT2: Enabling dense quantitative assessment of brain white matter connectivity using streamlines tractography. NeuroImage, 119, 338–351. https://doi.org/10.1016/j.neuroimage.2015.06.092

Smith, S. M., Jenkinson, M., Woolrich, M. W., Beckmann, C. F., Behrens, T. E. J., Johansen-Berg, H., Bannister, P. R., De Luca, M., Drobnjak, I., Flitney, D. E., Niazy, R. K., Saunders, J., Vickers, J., Zhang, Y., De Stefano, N., Brady, J. M., & Matthews, P. M. (2004). Advances in functional and structural MR image analysis and implementation as FSL. NeuroImage, 23, S208–S219. https://doi.org/10.1016/j.neuroimage.2004.07.051

Sotiropoulos, S. N., & Zalesky, A. (2019). Building connectomes using diffusion MRI: Why, how and but. NMR in Biomedicine, 32(4), e3752. https://doi.org/10.1002/nbm.3752

Sporns, O., Tononi, G., & Kötter, R. (2005). The Human Connectome: A Structural Description of the Human Brain. PLOS Computational Biology, 1(4), e42. https://doi.org/10.1371/journal.pcbi.0010042

Sun, L., Liang, X., Duan, D., Liu, J., Chen, Y., Wang, X., Liao, X., Xia, M., Zhao, T., & He, Y. (2022). Structural insight into the individual variability architecture of the functional brain connectome. NeuroImage, 259, 119387. https://doi.org/10.1016/j.neuroimage.2022.119387

Thomas, C., Ye, F. Q., Irfanoglu, M. O., Modi, P., Saleem, K. S., Leopold, D. A., & Pierpaoli, C. (2014). Anatomical accuracy of brain connections derived from diffusion MRI tractography is inherently limited. Proceedings of the National Academy of Sciences of the United States of America, 111(46), 16574–16579. https://doi.org/10.1073/pnas.1405672111

Tournier, J.-D., Calamante, F., & Connelly, A. (n.d.). Improved probabilistic streamlines tractography by 2nd order integration over fibre orientation distributions. 1.

Tournier, J.-D., Calamante, F., & Connelly, A. (2007). Robust determination of the fibre orientation distribution in diffusion MRI: Non-negativity constrained super-resolved spherical deconvolution. NeuroImage, 35(4), 1459–1472. https://doi.org/10.1016/j.neuroimage.2007.02.016

Tournier, J.-D., Calamante, F., & Connelly, A. (2012). MRtrix: Diffusion tractography in crossing fiber regions. International Journal of Imaging Systems and Technology, 22(1), 53–66. https://doi.org/10.1002/ima.22005

Tournier, J.-D., Smith, R., Raffelt, D., Tabbara, R., Dhollander, T., Pietsch, M., Christiaens, D., Jeurissen, B., Yeh, C.-H., & Connelly, A. (2019). MRtrix3: A fast, flexible and open software framework for medical image processing and visualisation. NeuroImage, 202, 116137. https://doi.org/10.1016/j.neuroimage.2019.116137

van den Heuvel, M. P., Kahn, R. S., Goñi, J., & Sporns, O. (2012). High-cost, high-capacity backbone for global brain communication. Proceedings of the National Academy of Sciences, 109(28), 11372–11377. https://doi.org/10.1073/pnas.1203593109

van den Heuvel, M. P., & Sporns, O. (2013a). An anatomical substrate for integration among functional networks in human cortex. The Journal of Neuroscience: The Official Journal of the Society for Neuroscience, 33(36), 14489–14500. https://doi.org/10.1523/JNEUROSCI.2128-13.2013

van den Heuvel, M. P., & Sporns, O. (2013b). Network hubs in the human brain. Trends in Cognitive Sciences, 17(12), 683–696. https://doi.org/10.1016/j.tics.2013.09.012

Van Essen, D. C., Glasser, M. F., Dierker, D. L., Harwell, J., & Coalson, T. (2012). Parcellations and Hemispheric Asymmetries of Human Cerebral Cortex Analyzed on Surface-Based Atlases. Cerebral Cortex, 22(10), 2241–2262. https://doi.org/10.1093/cercor/bhr291

Varshney, L. R., Chen, B. L., Paniagua, E., Hall, D. H., & Chklovskii, D. B. (2011). Structural Properties of the Caenorhabditis elegans Neuronal Network. PLOS Computational Biology, 7(2), e1001066. https://doi.org/10.1371/journal.pcbi.1001066

Wang, C., Yoldemir, B., & Abugharbieh, R. (2015). Multimodal Cortical Parcellation Based on Anatomical and Functional Brain Connectivity. In N. Navab, J. Hornegger, W. M. Wells, & A. F. Frangi (Eds.), Medical Image Computing and Computer-Assisted Intervention – MICCAI 2015 (pp. 21– 28). Springer International Publishing. https://doi.org/10.1007/978-3-319-24574-4_3

Westfall, P. H. (2014). Kurtosis as Peakedness, 1905–2014. R.I.P. The American Statistician, 68(3), 191–195. https://doi.org/10.1080/00031305.2014.917055

Yan, X., Kong, R., Xue, A., Yang, Q., Orban, C., An, L., Holmes, A. J., Qian, X., Chen, J., Zuo, X.-N., Zhou, J. H., Fortier, M. V., Tan, A. P., Gluckman, P., Chong, Y. S., Meaney, M., Bzdok, D., Eickhoff, S. B., & Yeo, B. T. T. (2022). Homotopic local-global parcellation of the human cerebral cortex from resting-state functional connectivity (p. 2022.10.25.513788). bioRxiv. https://doi.org/10.1101/2022.10.25.513788

Yeh, F.-C., Vettel, J. M., Singh, A., Poczos, B., Grafton, S. T., Erickson, K. I., Tseng, W.-Y. I., & Verstynen, T. D. (2016). Quantifying Differences and Similarities in Whole-Brain White Matter Architecture Using Local Connectome Fingerprints. PLOS Computational Biology, 12(11), e1005203. https://doi.org/10.1371/journal.pcbi.1005203

Zalesky, A., Fornito, A., Harding, I. H., Cocchi, L., Yücel, M., Pantelis, C., & Bullmore, E. T. (2010). Whole-brain anatomical networks: Does the choice of nodes matter? NeuroImage, 50(3), 970–983. https://doi.org/10.1016/j.neuroimage.2009.12.027

Zhang, Y., Brady, M., & Smith, S. (2001). Segmentation of brain MR images through a hidden Markov random field model and the expectation-maximization algorithm. IEEE Transactions on Medical Imaging, 20(1), 45–57. https://doi.org/10.1109/42.906424

Zilles, K. (2018). Brodmann: A pioneer of human brain mapping—his impact on concepts of cortical organization. Brain, 141(11), 3262–3278. https://doi.org/10.1093/brain/awy273

Zucca, R., Arsiwalla, X. D., Le, H., Rubinov, M., Gurguí, A., & Verschure, P. (2019). The Degree Distribution of Human Brain Functional Connectivity is Generalized Pareto: A Multi-Scale Analysis (p. 840066). bioRxiv. https://doi.org/10.1101/840066

