## Supplementary Figures for "Can hubs of the human connectome be identified consistently with diffusion MRI?"

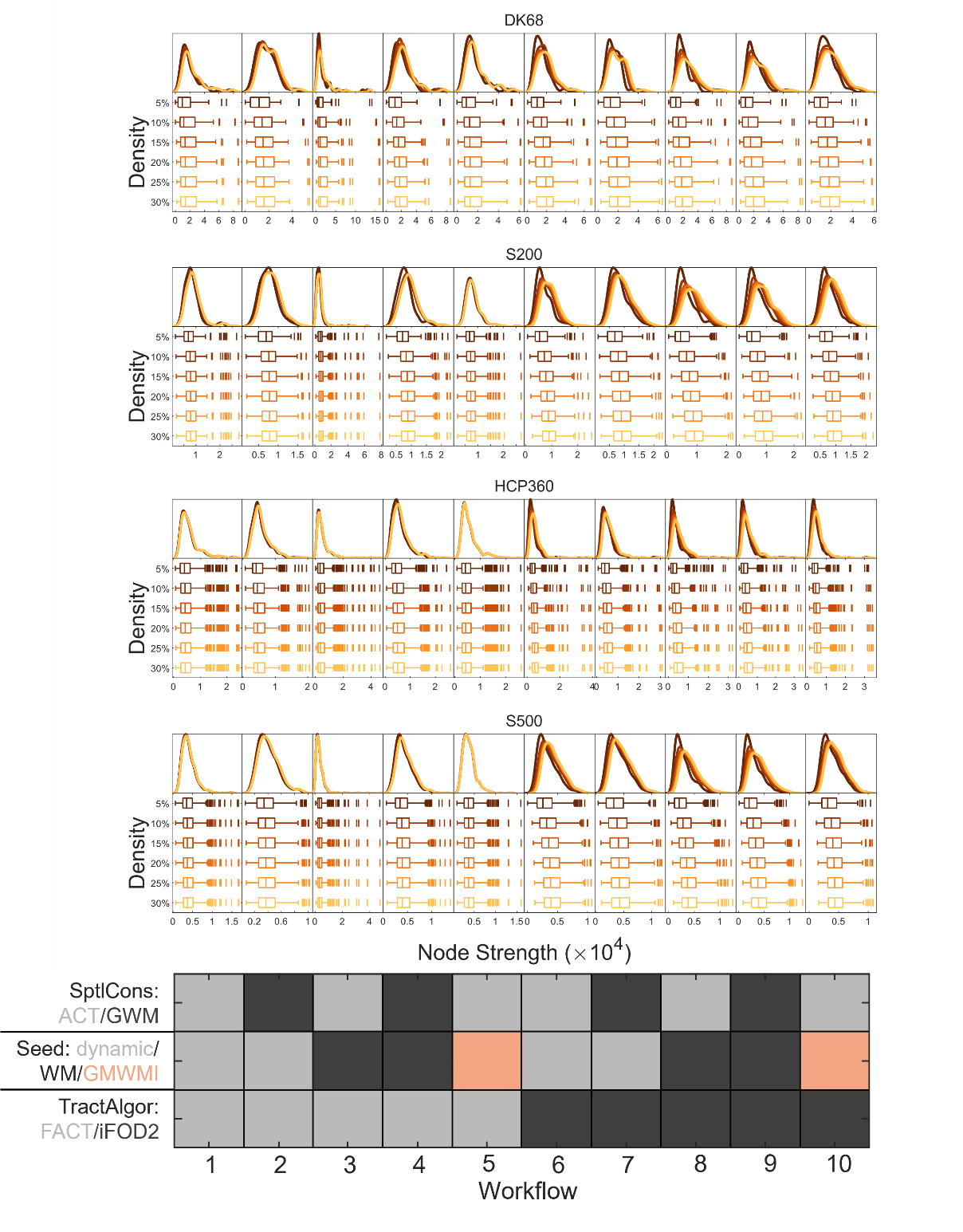

**Figure S1: Effect of threshold density on strength distributions across parcellations and workflows** (edges retained using edge coefficient-of-variation). Across all parcellations, workflows 1, 3, 5 (ACT+FACT) tend to have more outliers (more than 1.5 times the interquartile range above the third quartile, or more than 1.5 times the interquartile range below the first quartile), whilst workflows 6-10 (iFOD2) are more affected by changes in density.

Parcellation: DK68 = Desikan-Killiany 68 nodes, S200 = Schaefer 200 nodes, HCP360 = Glasser 360 nodes, S500 = Schaefer 500 nodes. Tractography: SptlCons = spatial constraints on streamline propagation, ACT = anatomically constrained tractography, GWM = grey-white masking; Seed = streamline seeding algorithm, dynamic = dynamic seeding, WM = white matter seeding, GMWMI = grey matter-white matter interface seeding; TractAlgor = Streamline tractography algorithm, FACT = fiber assignment by continuous tractography, iFOD2 = second-order integration over fiber orientation distributions.

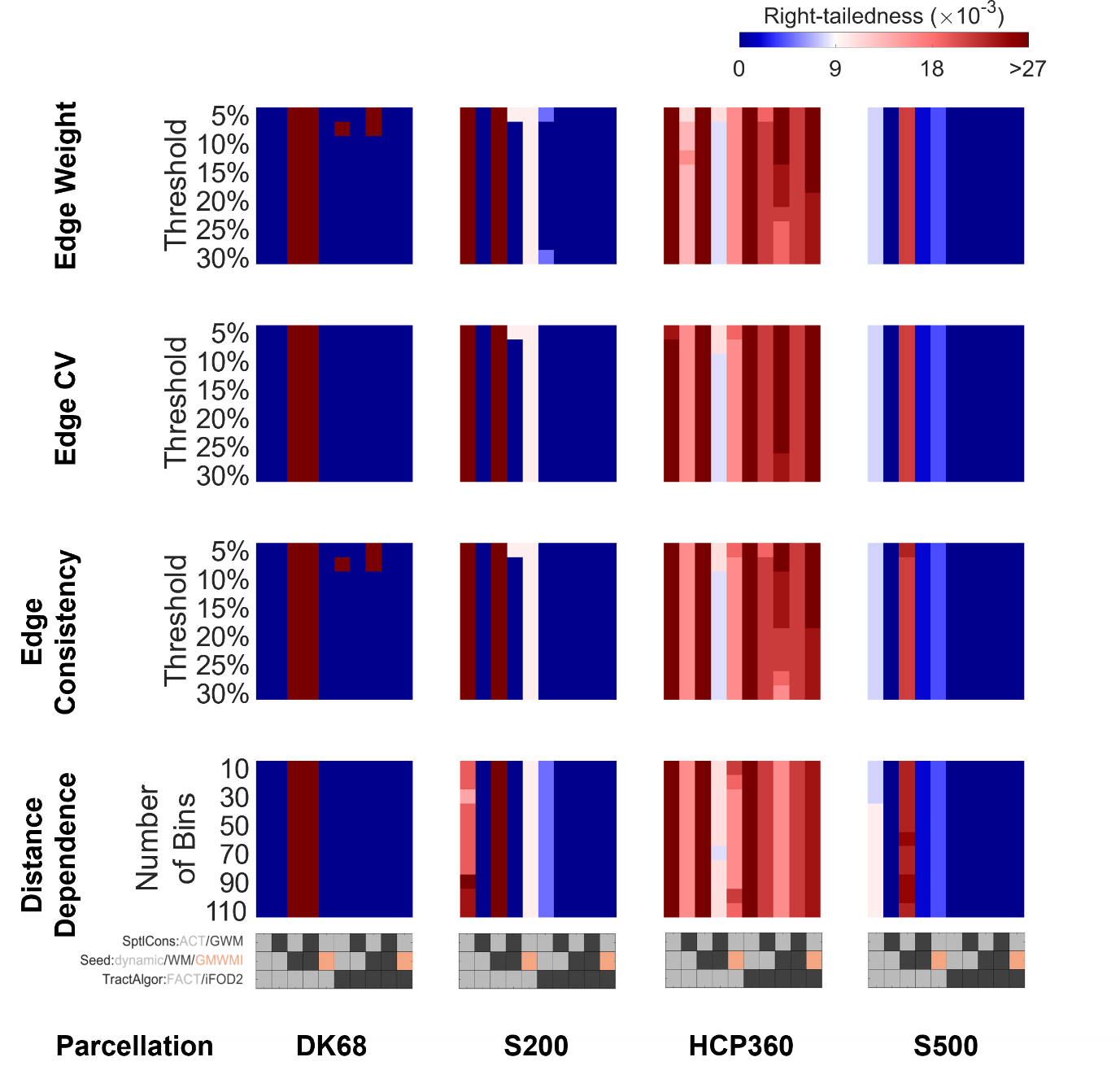

**Figure S2: Comparison of the right-tailedness of strength distributions across all parcellations, tractography workflows, group reconstructions, and thresholds** (see Methods for calculation of right-tailedness). The parcellation and workflow vary along the x-axis, and the group reconstruction and threshold vary along the y-axis. Each subplot shows the right-tailedness as a function of tractography workflow and threshold for a given parcellation and group reconstruction. The range of cool/warm colors correspond to a right-tailedness less than/greater than that of the exponential distribution.

Parcellation: DK68 = Desikan-Killiany 68 nodes, S200 = Schaefer 200 nodes, HCP360 = Glasser 360 nodes, S500 = Schaefer 500 nodes. Tractography: SptlCons = spatial constraints on streamline propagation, ACT = anatomically constrained tractography, GWM = grey-white masking; Seed = streamline seeding algorithm, dynamic = dynamic seeding, WM = white matter seeding, GMWMI = grey matter-white matter interface seeding; TractAlgor = Streamline tractography algorithm, FACT = fiber assignment by continuous tractography, iFOD2 = second-order integration over fiber orientation distributions.

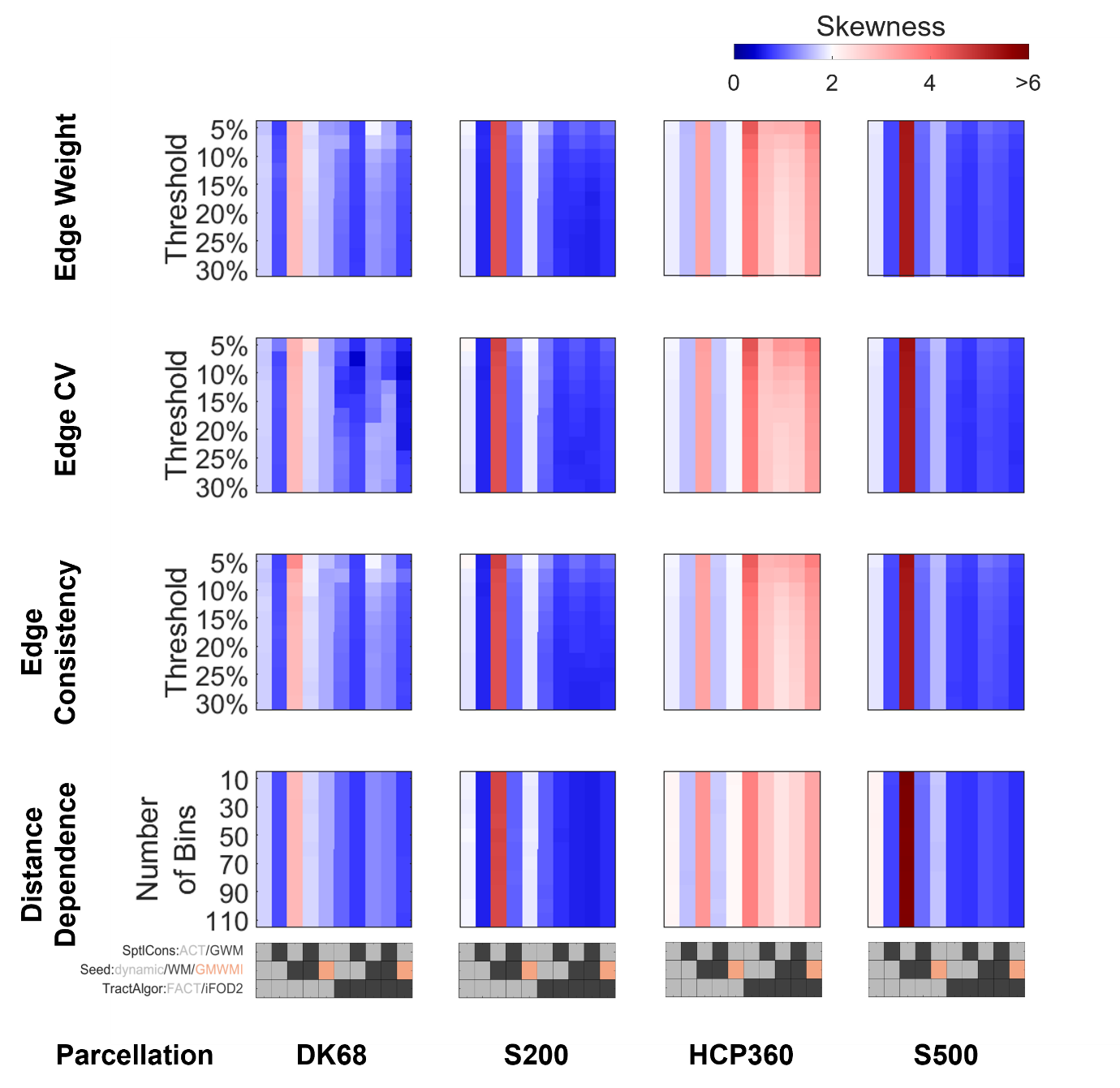

**Figure S3: Comparison of the skewness of strength distributions across all parcellations, tractography workflows, group reconstructions, and thresholds**. The parcellation and workflow vary along the x-axis, and the group reconstruction and threshold vary along the y-axis. Each subplot shows the skewness as a function of tractography workflow and threshold for a given parcellation and group reconstruction. The range of cool/warm colors correspond to a skewness less than/greater than that of the exponential distribution.

Parcellation: DK68 = Desikan-Killiany 68 nodes, S200 = Schaefer 200 nodes, HCP360 = Glasser 360 nodes, S500 = Schaefer 500 nodes. Tractography: SptlCons = spatial constraints on streamline propagation, ACT = anatomically constrained tractography, GWM = grey-white masking; Seed = streamline seeding algorithm, dynamic = dynamic seeding, WM = white matter seeding, GMWMI = grey matter-white matter interface seeding; TractAlgor = Streamline tractography algorithm, FACT = fiber assignment by continuous tractography, iFOD2 = second-order integration over fiber orientation distributions.

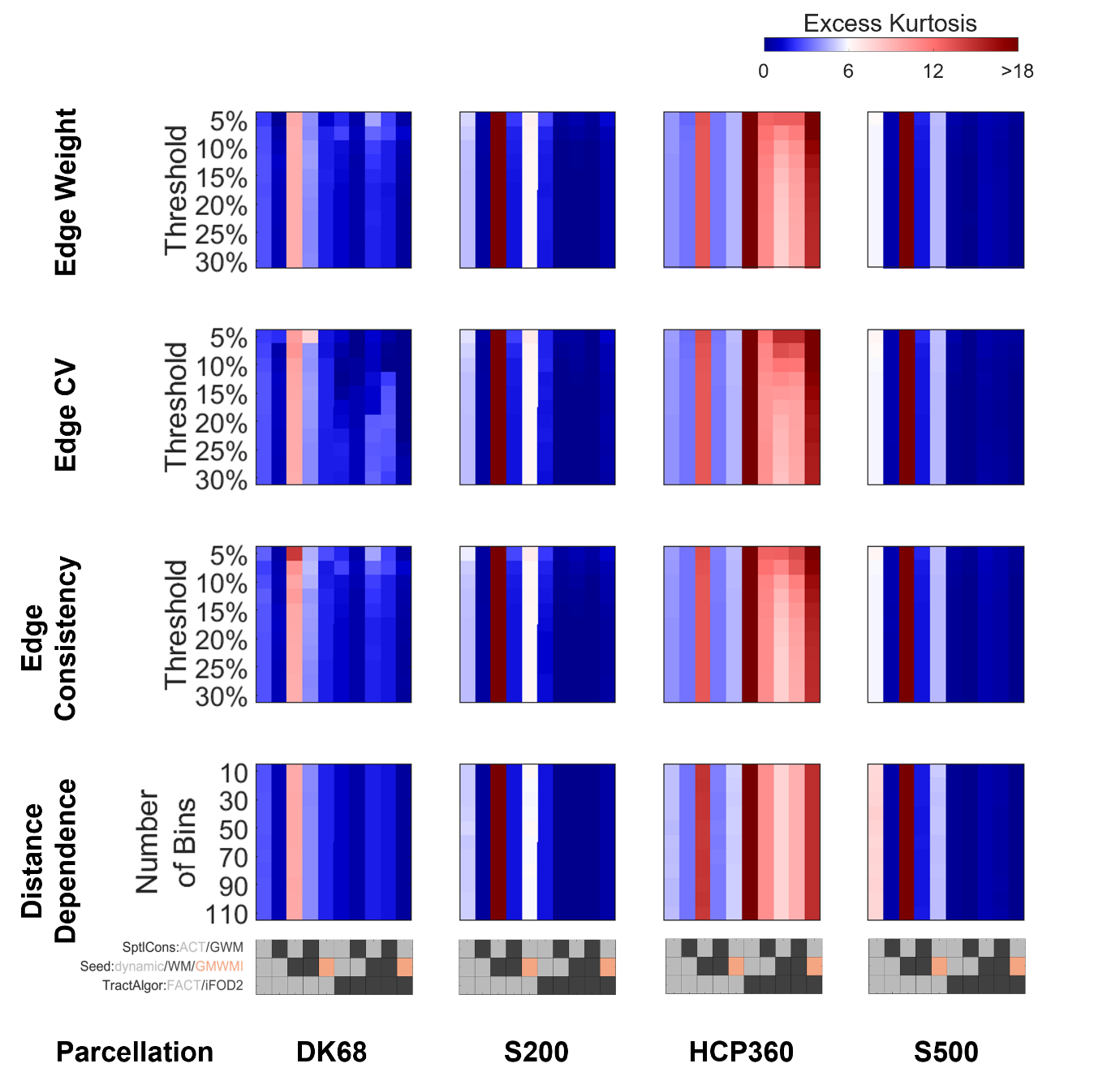

**Figure S4: Comparison of the kurtosis of strength distributions across all parcellations, tractography workflows, group reconstructions, and thresholds**. The parcellation and workflow vary along the x-axis, and the group reconstruction and threshold vary along the y-axis. Each subplot shows the kurtosis as a function of tractography workflow and threshold for a given parcellation and group reconstruction. The range of cool/warm colors correspond to a kurtosis less than/greater than that of the exponential distribution.

Parcellation: DK68 = Desikan-Killiany 68 nodes, S200 = Schaefer 200 nodes, HCP360 = Glasser 360 nodes, S500 = Schaefer 500 nodes. Tractography: SptlCons = spatial constraints on streamline propagation, ACT = anatomically constrained tractography, GWM = grey-white masking; Seed = streamline seeding algorithm, dynamic = dynamic seeding, WM = white matter seeding, GMWMI = grey matter-white matter interface seeding; TractAlgor = Streamline tractography algorithm, FACT = fiber assignment by continuous tractography, iFOD2 = second-order integration over fiber orientation distributions.

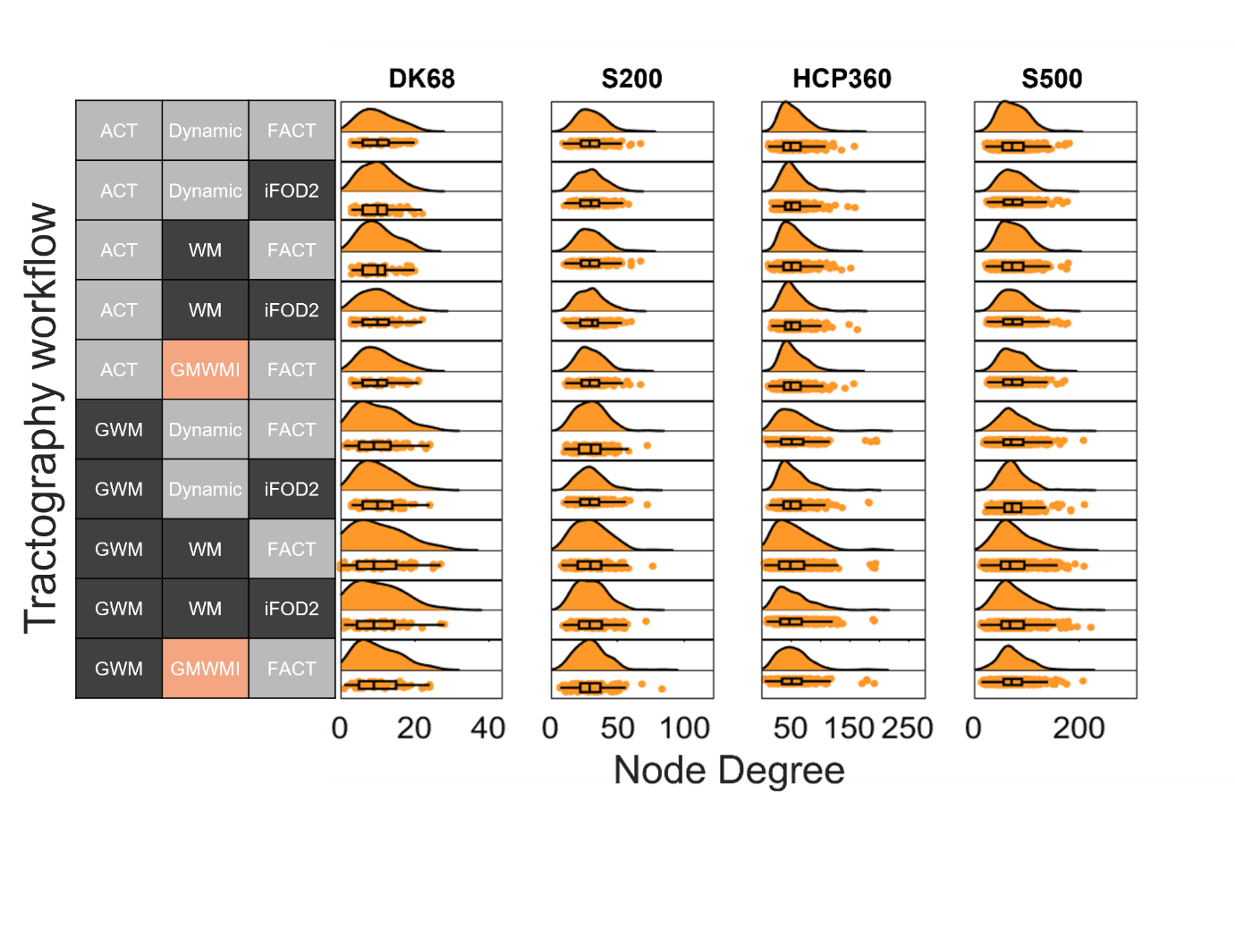

**Figure S5: Degree distributions of binarized connectomes**. The group connectomes are constructed using the edge coefficient-of-variation (CV) for each parcellation and workflow at a connection density of 20%.

Parcellation: DK68 = Desikan-Killiany 68 nodes, S200 = Schaefer 200 nodes, HCP360 = Glasser 360 nodes, S500 = Schaefer 500 nodes. Tractography: SptlCons = spatial constraints on streamline propagation, ACT = anatomically constrained tractography, GWM = grey-white masking; Seed = streamline seeding algorithm, dynamic = dynamic seeding, WM = white matter seeding, GMWMI = grey matter-white matter interface seeding; TractAlgor = Streamline tractography algorithm, FACT = fiber assignment by continuous tractography, iFOD2 = second-order integration over fiber orientation distributions.

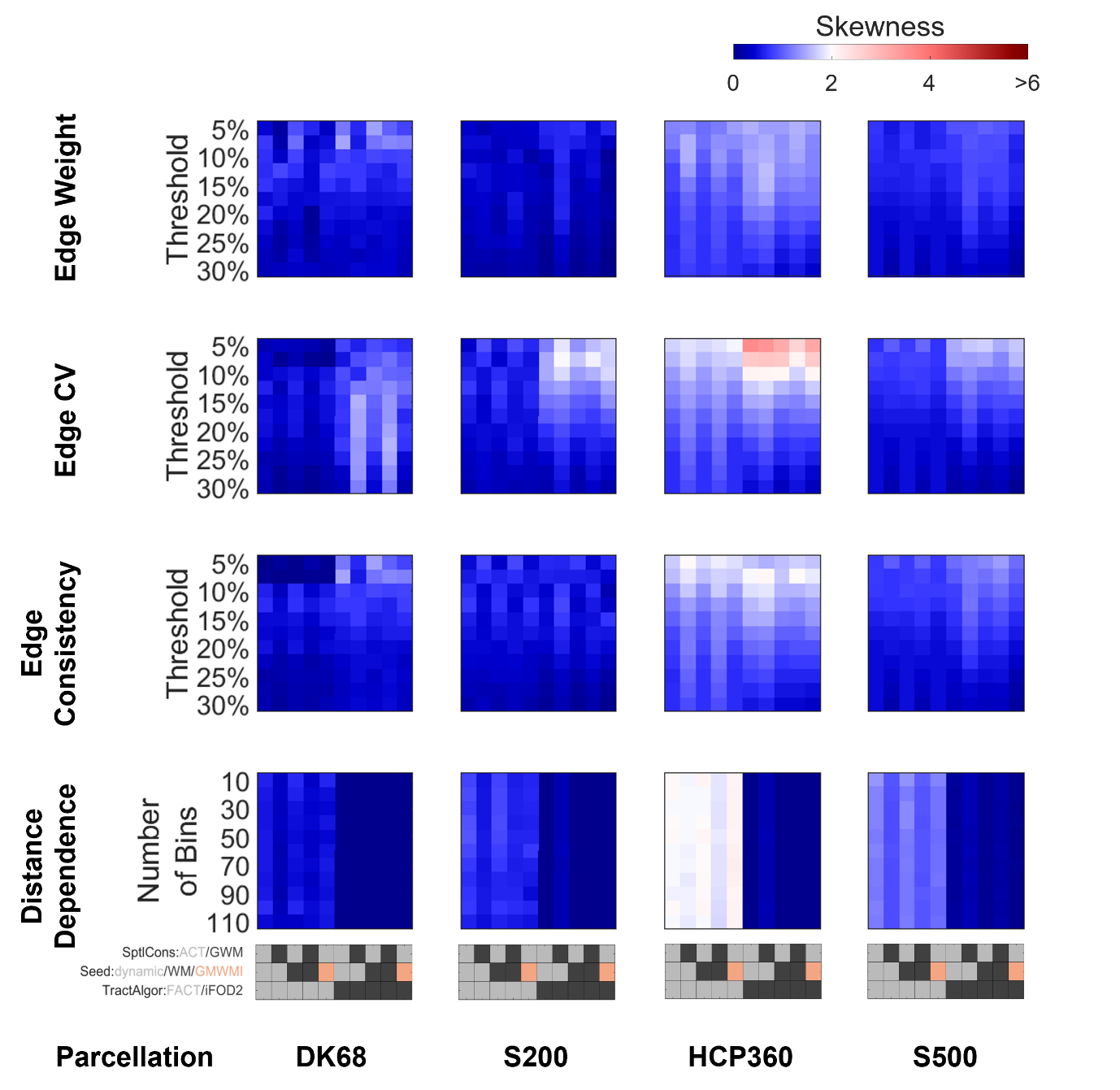

**Figure S6: Comparison of skewness of degree distribution across all parcellations, tractography workflows, group reconstructions, and thresholds in binarized connectomes**. The parcellation and workflow vary along the x-axis, and the group reconstruction and threshold vary along the y-axis. Each subplot shows the skewness as a function of tractography workflow and threshold for a given parcellation and group reconstruction. The range of cool/warm colors correspond to a skewness less than/greater than that of the exponential distribution.

Parcellation: DK68 = Desikan-Killiany 68 nodes, S200 = Schaefer 200 nodes, HCP360 = Glasser 360 nodes, S500 = Schaefer 500 nodes. Tractography: SptlCons = spatial constraints on streamline propagation, ACT = anatomically constrained tractography, GWM = grey-white masking; Seed = streamline seeding algorithm, dynamic = dynamic seeding, WM = white matter seeding, GMWMI = grey matter-white matter interface seeding; TractAlgor = Streamline tractography algorithm, FACT = fiber assignment by continuous tractography, iFOD2 = second-order integration over fiber orientation distributions.

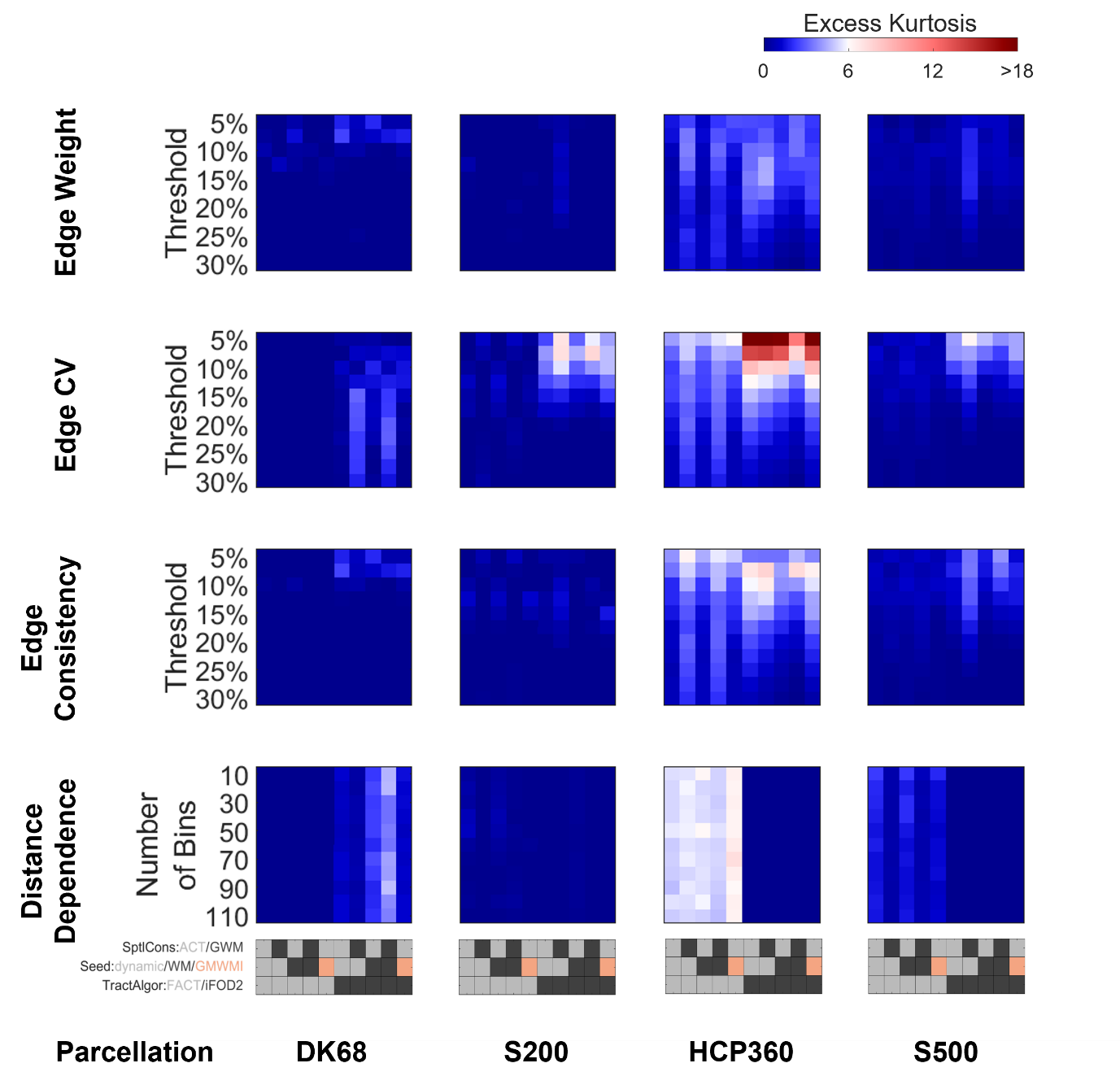

**Figure S7: Comparison of kurtosis of degree distribution across all parcellations, tractography workflows, group reconstructions, and thresholds in binarized connectomes**. The parcellation and workflow vary along the x-axis, and the group reconstruction and threshold vary along the y-axis. Each subplot shows the kurtosis as a function of tractography workflow and threshold for a given parcellation and group reconstruction. The range of cool/warm colors correspond to a kurtosis less than/greater than that of the exponential distribution.

Parcellation: DK68 = Desikan-Killiany 68 nodes, S200 = Schaefer 200 nodes, HCP360 = Glasser 360 nodes, S500 = Schaefer 500 nodes. Tractography: SptlCons = spatial constraints on streamline propagation, ACT = anatomically constrained tractography, GWM = grey-white masking; Seed = streamline seeding algorithm, dynamic = dynamic seeding, WM = white matter seeding, GMWMI = grey matter-white matter interface seeding; TractAlgor = Streamline tractography algorithm, FACT = fiber assignment by continuous tractography, iFOD2 = second-order integration over fiber orientation distributions.

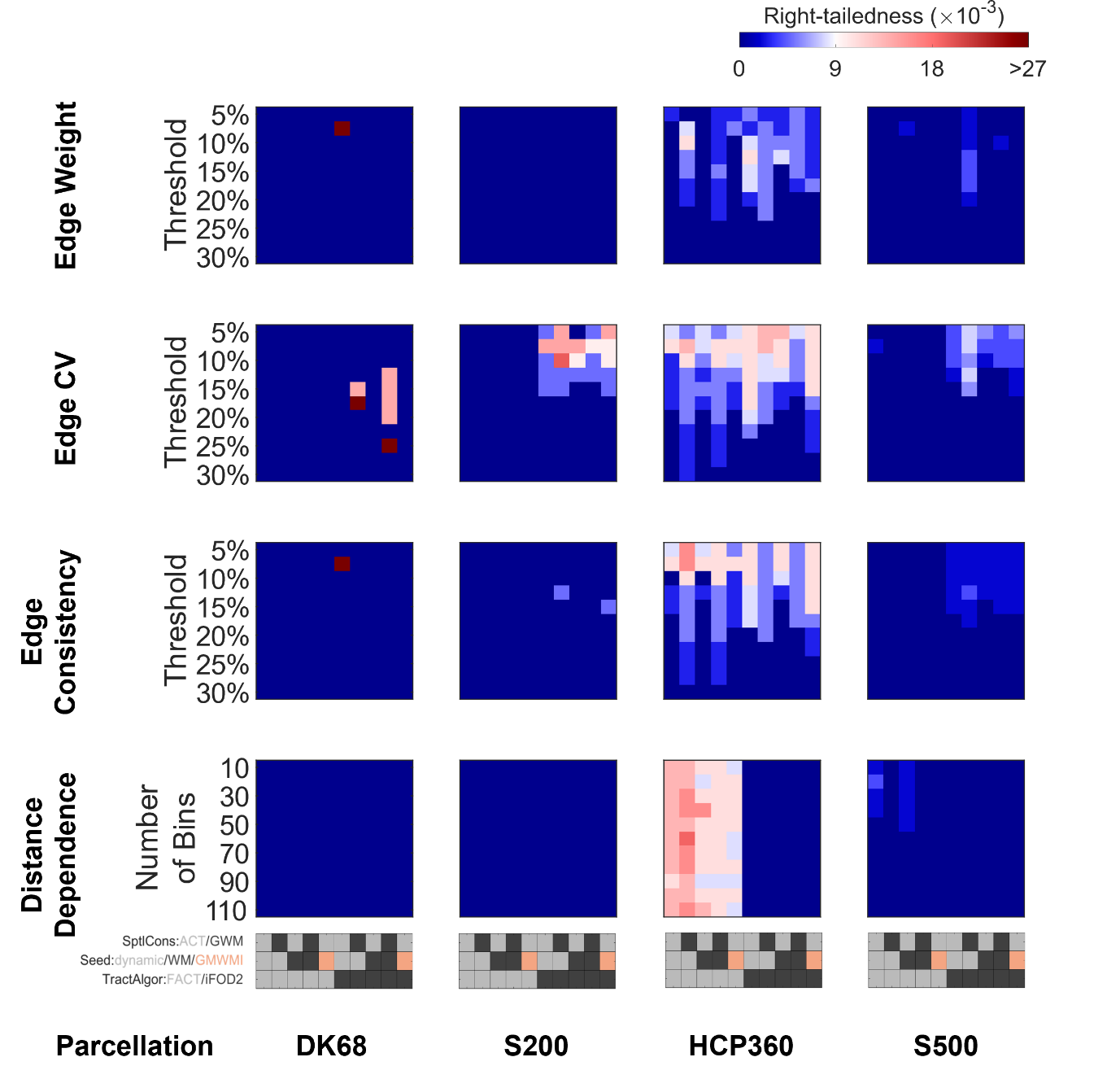

**Figure S8: Comparison of right-tailedness of strength distribution across all parcellations, tractography workflows, group reconstructions, and thresholds in binarized connectomes** (see Methods for calculation of right-tailedness). The parcellation and workflow vary along the x-axis, and the group reconstruction and threshold vary along the y-axis. Each subplot shows the right-tailedness as a function of tractography workflow and threshold for a given parcellation and group reconstruction. The range of cool/warm colours correspond to a right-tailedness less than/greater than that of the exponential distribution.

Parcellation: DK68 = Desikan-Killiany 68 nodes, S200 = Schaefer 200 nodes, HCP360 = Glasser 360 nodes, S500 = Schaefer 500 nodes. Tractography: SptlCons = spatial constraints on streamline propagation, ACT = anatomically constrained tractography, GWM = grey-white masking; Seed = streamline seeding algorithm, dynamic = dynamic seeding, WM = white matter seeding, GMWMI = grey matter-white matter interface seeding; TractAlgor = Streamline tractography algorithm, FACT = fiber assignment by continuous tractography, iFOD2 = second-order integration over fiber orientation distributions.

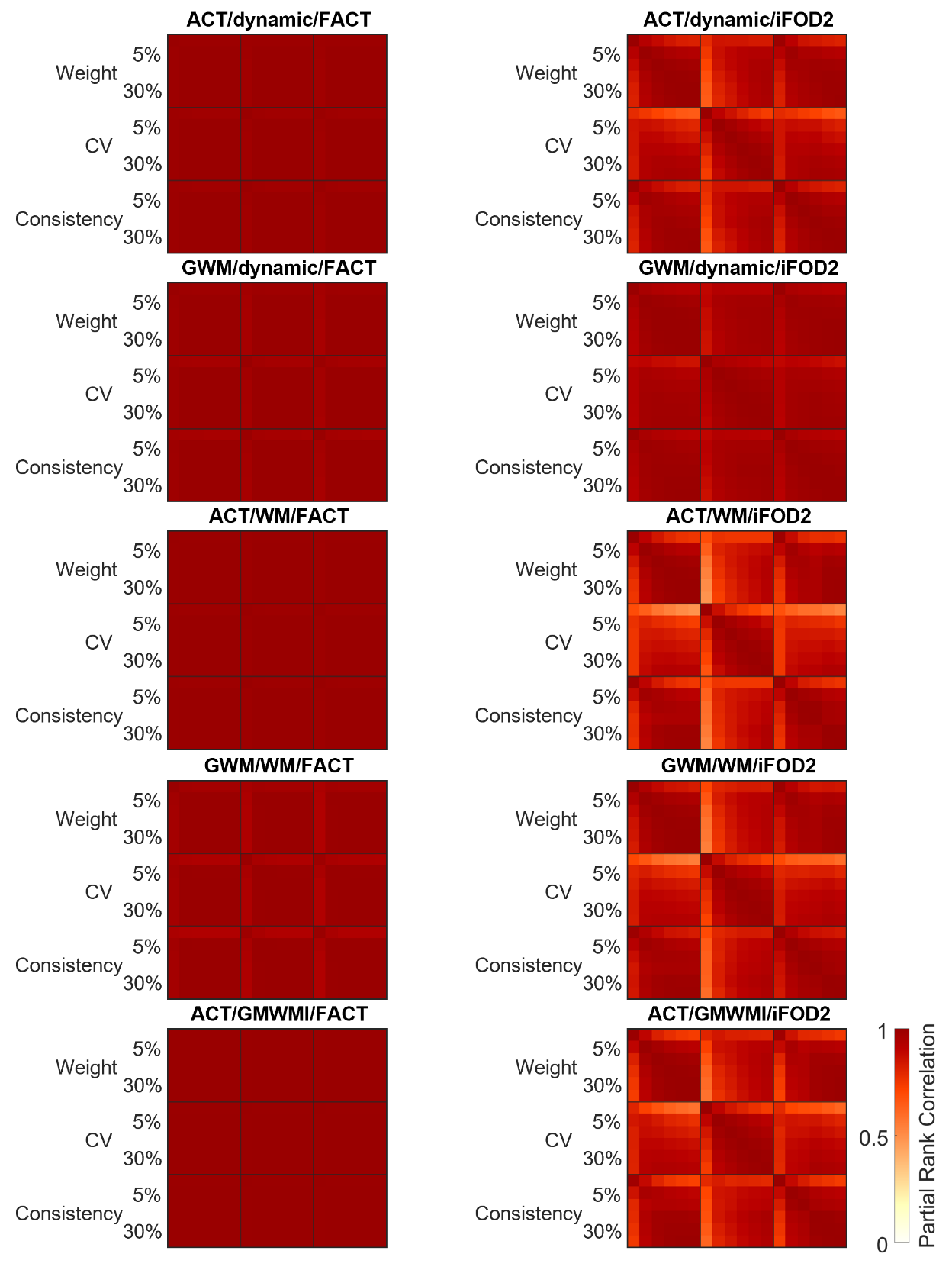

**Figure S9: Comparison of similarities between tractography, edge reconstruction, and density using the S200 parcellation in weighted connectomes**. Each plot represents the partial rank correlation (when corrected for surface area) between edge reconstruction parameters (edge weight, coefficient-of-variation, or consistency) and density (5, 10, 15, 20, 25, or 30%) for the ten different workflows assessed. Workflows are extremely consistent when using deterministic tractography (the minimum value of Spearman’s ρ in the first column is 0.93). When using probabilistic tractography (second column), the lowest values of Spearman’s ρ (0.64, 0.85, 0.49, 0.54, 0.58) are all found when comparing 5% threshold to 30% threshold.

ACT = anatomically constrained tractography, GWM = grey-white masking; dynamic = dynamic seeding, WM = white matter seeding, GMWMI = grey matter-white matter interface seeding; FACT = fiber assignment by continuous tractography, iFOD2 = second-order integration over fiber orientation distributions.

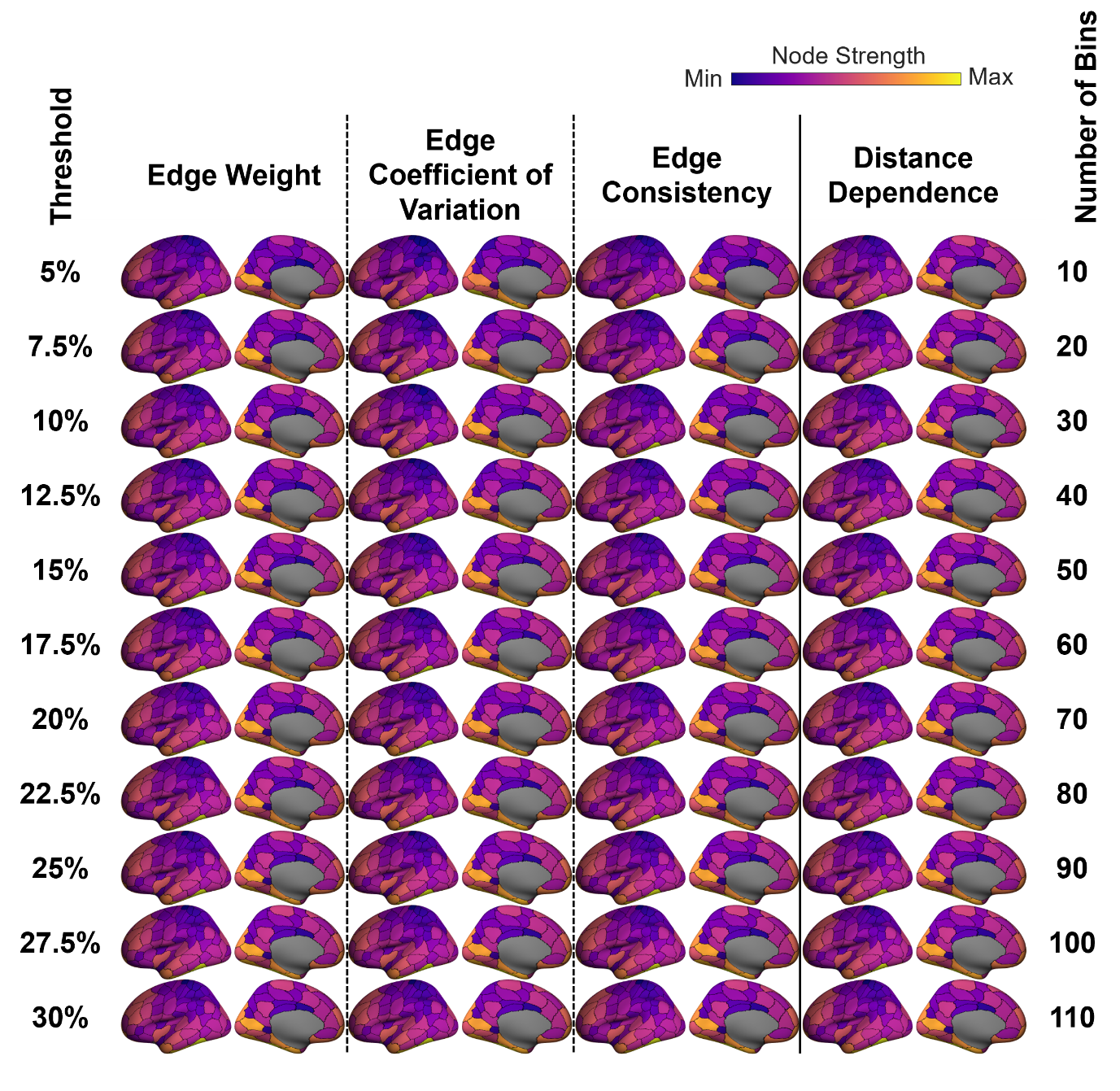

**Figure S10: Spatial maps of node strengths for different group reconstructions and thresholds**. The first 3 columns use algorithms that retain the top T% of edges with a specified property (edge weight = mean across participants; coefficient of variation = standard deviation ÷ mean across participants; consistency = fraction of participants where the edge is present), where T is the threshold percentage. The rightmost column used an algorithm described in (Betzel et al., 2019) where the number of bins, rather than a density threshold, is used to permute edges (see Methods). Here, the S200 parcellation and tractography workflow #7 (GWM/dynamic seeding/iFOD2) were used.

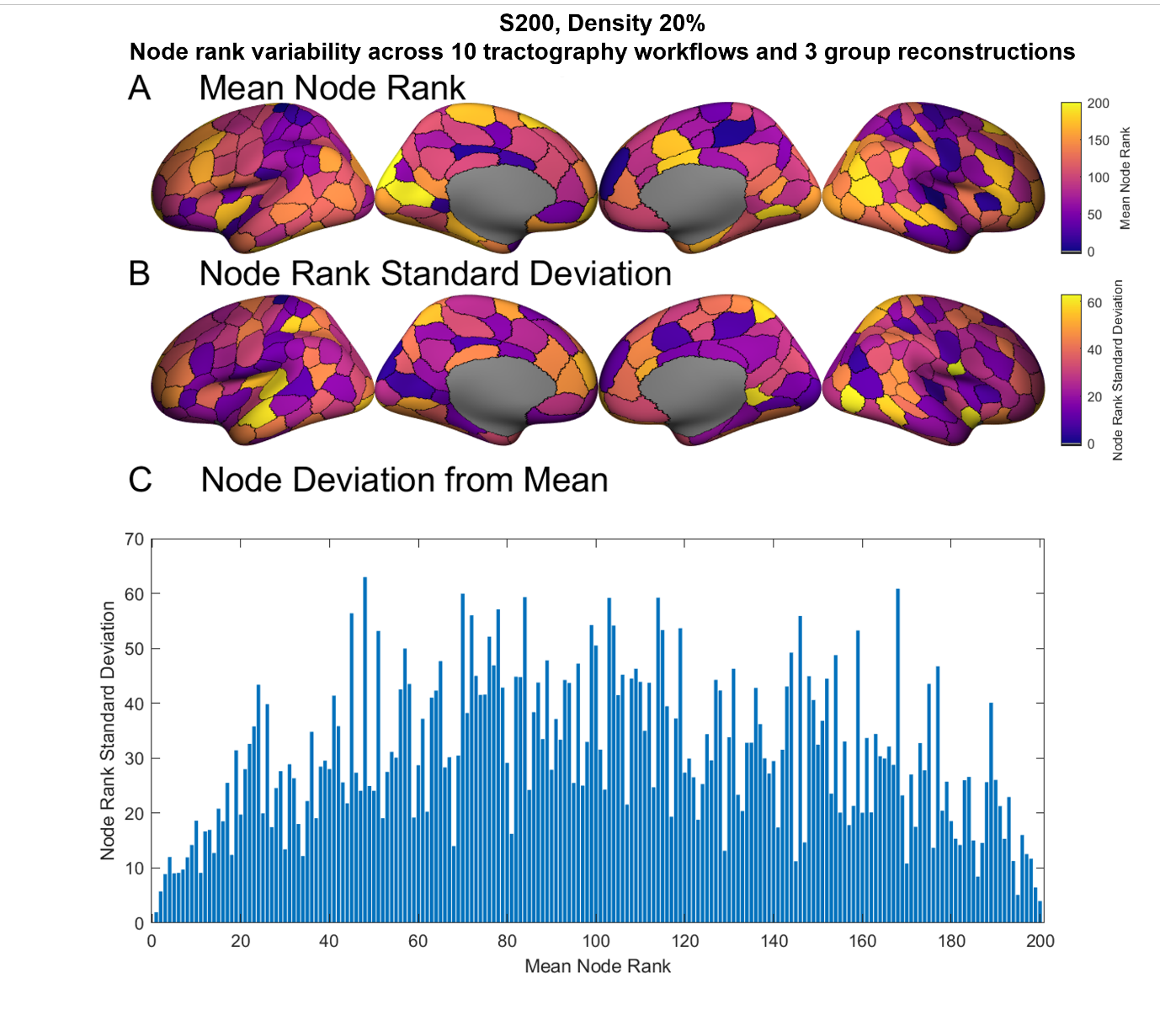

**Figure S11: Node rank variability in an example parcellation and density as tractography workflow and group reconstruction are varied.** Here, the S200 parcellation and a density of 20% are used, the same as the example in Figure 5 left. Node ranks across all tractography workflows and density-matched group reconstructions (weight, consistency, and coefficient-of-variation) were calculated. (A) Mean node ranks for across the combinations of workflows and reconstructions. Note that purple colours represent more strongly connected nodes. (B) The standard deviation of node rank across the combinations of workflows and reconstructions. Note that purple colours represent more nodes with less variable rankings. (C) Standard deviations of each node from its mean ranking. Nodes are sorted in order of their mean rank.

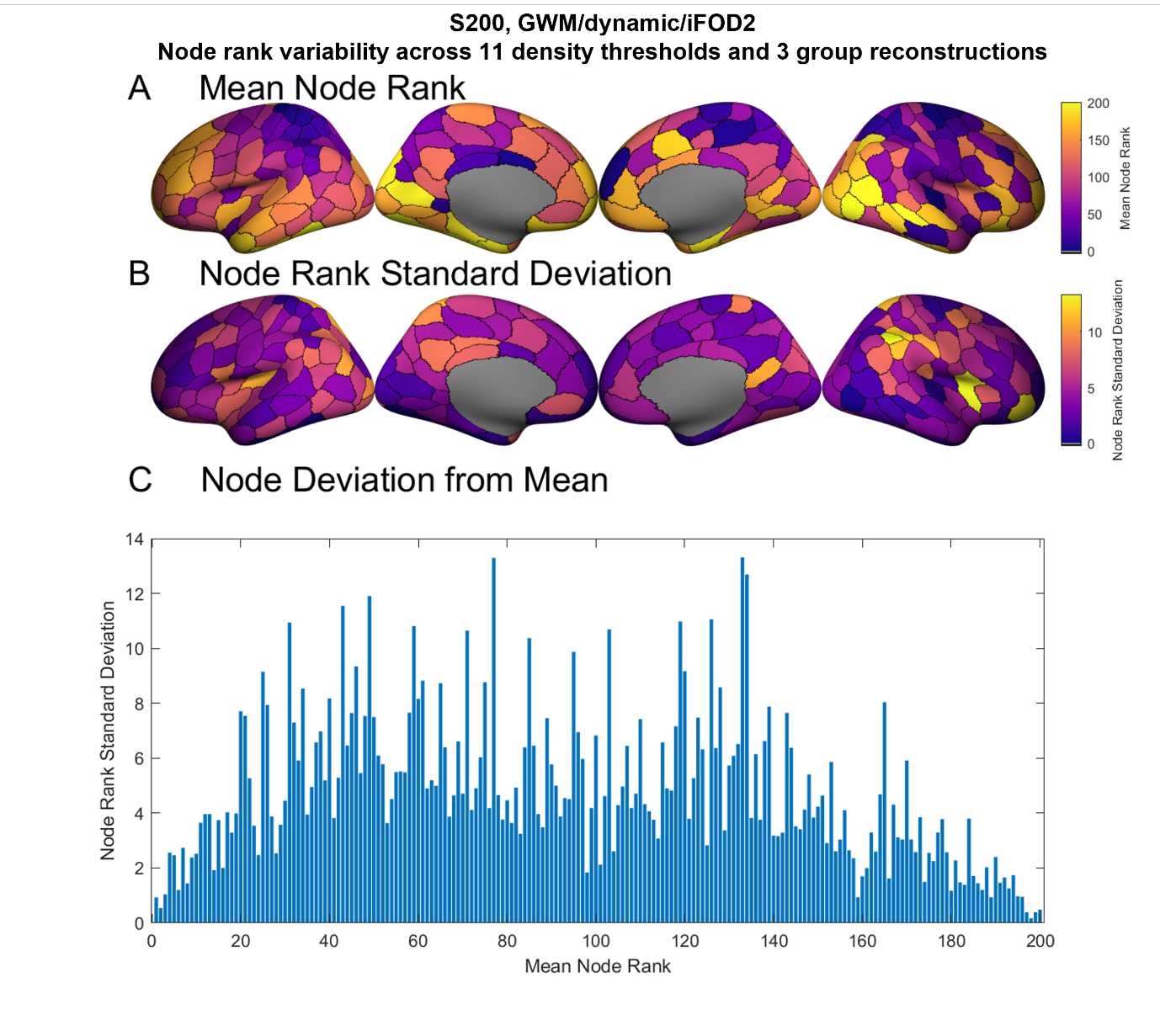

**Figure S12: Node rank variability in an example parcellation and workflow as group reconstruction and density are varied.** Here, the S200 parcellation and workflow 7 (GWM/dynamic/iFOD2) are used, the same as the example in Figure 5 left. Node ranks across all density thresholds (5% - 30%) and density-matched group reconstructions (weight, consistency, and coefficient-of-variation) were calculated. (A) Mean node ranks for across the combinations of group reconstructions and densities. Note that purple colours represent more strongly connected nodes. (B) The standard deviation of node rank across the combinations of group reconstructions and densities. Note that purple colours represent more nodes with less variable rankings. (C) Standard deviations of each node from its mean ranking. Nodes are sorted in order of their mean rank.

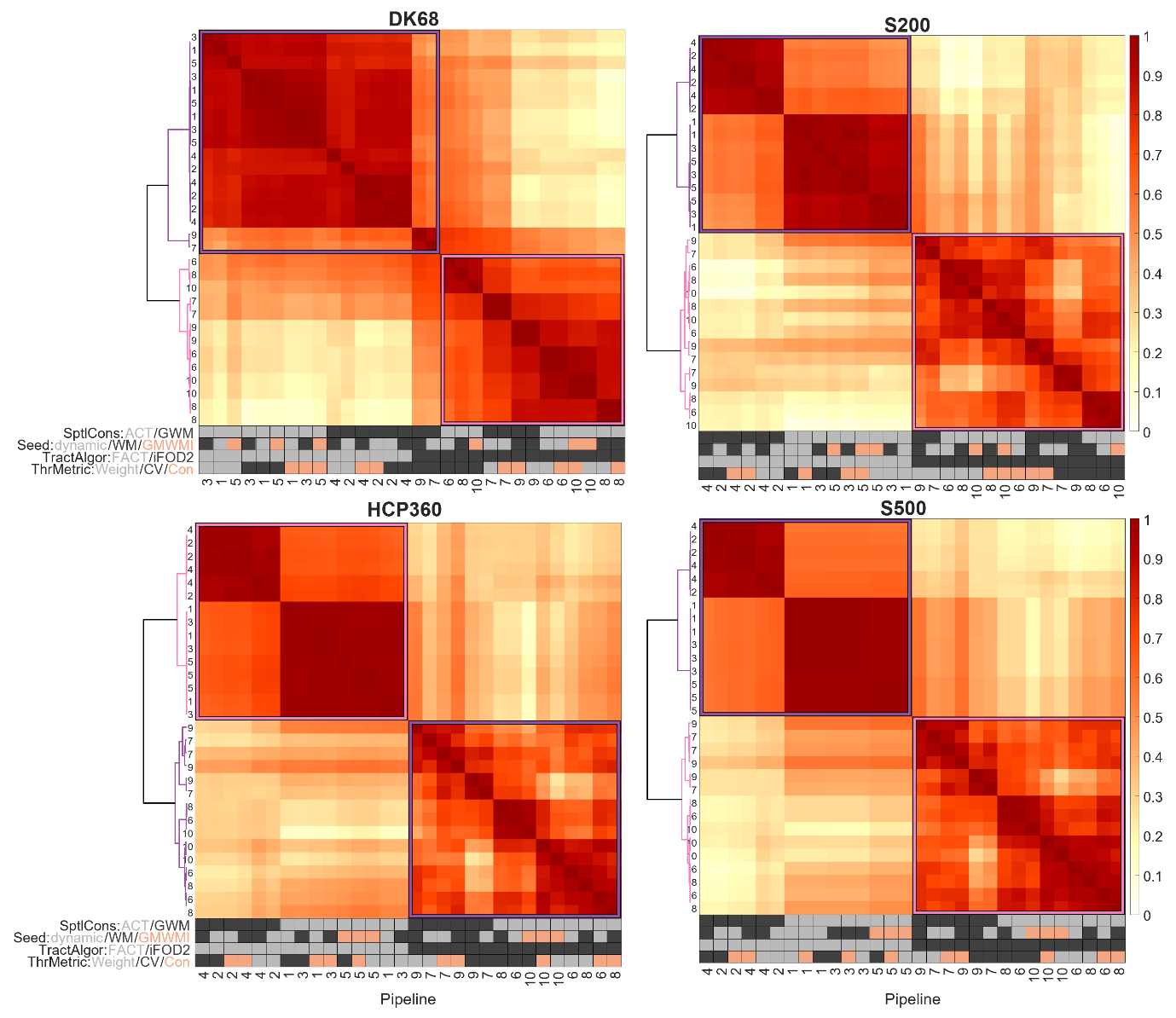

**Figure S13: Similarity between tractography workflows and group reconstruction metrics for each parcellation using binarized connectomes with density 20%.** Each heatmap shows partial rank correlations of node degree in binarized connectomes, corrected for surface area. Workflows are reordered using hierarchical clustering.

Parcellation: DK68 = Desikan-Killiany 68 nodes, S200 = Schaefer 200 nodes, HCP360 = Glasser 360 nodes, S500 = Schaefer 500 nodes. Tractography: SptlCons = spatial constraints on streamline propagation, ACT = anatomically constrained tractography, GWM = grey-white masking; Seed = streamline seeding algorithm, dynamic = dynamic seeding, WM = white matter seeding, GMWMI = grey matter-white matter interface seeding; TractAlgor = Streamline tractography algorithm, FACT = fiber assignment by continuous tractography, iFOD2 = second-order integration over fiber orientation distributions. Group aggregation: ThrMetric = group-reconstruction thresholding metric, Weight = edge weight, CV = edge coefficient-of-variation, Con = edge consistency.

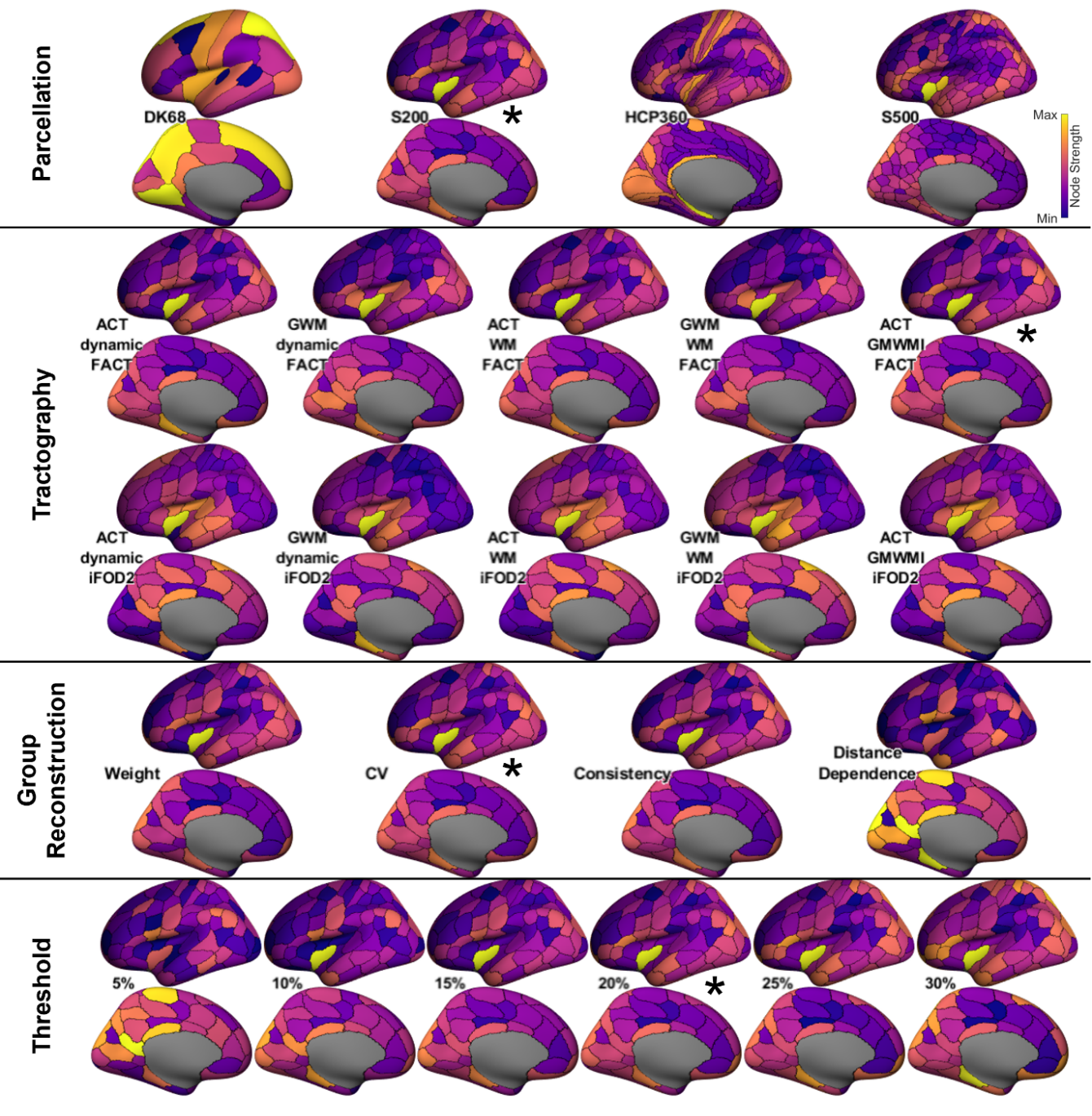

**Figure S14: Spatial maps of nodes degrees using different processing workflows in binarized connectomes**. Cortical parcellation, tractography parameters, group reconstruction, and density threshold were independently compared. Here, the default parameters are indicated by stars: parcellation, Schaefer 200; workflow, ACT/GMWMI/FACT; group reconstruction, edge CV; density threshold, 20%. Each row independently alters one option. Lateral and medial views of only the left hemisphere are presented for brevity. Each image is independently colored. Asterisk denotes the parameter value used throughout other groups, i.e., the starred images are the same.

Parcellation: DK68 = Desikan-Killiany 68 nodes, S200 = Schaefer 200 nodes, HCP360 = Glasser 360 nodes, S500 = Schaefer 500 nodes. Tractography: ACT = anatomically constrained tractography, GWM = grey-white masking; dynamic = dynamic seeding, WM = white matter seeding, GMWMI = grey matter-white matter interface seeding; FACT = fiber assignment by continuous tractography, iFOD2 = second-order integration over fiber orientation distributions. Group reconstruction: CV = coefficient of variation
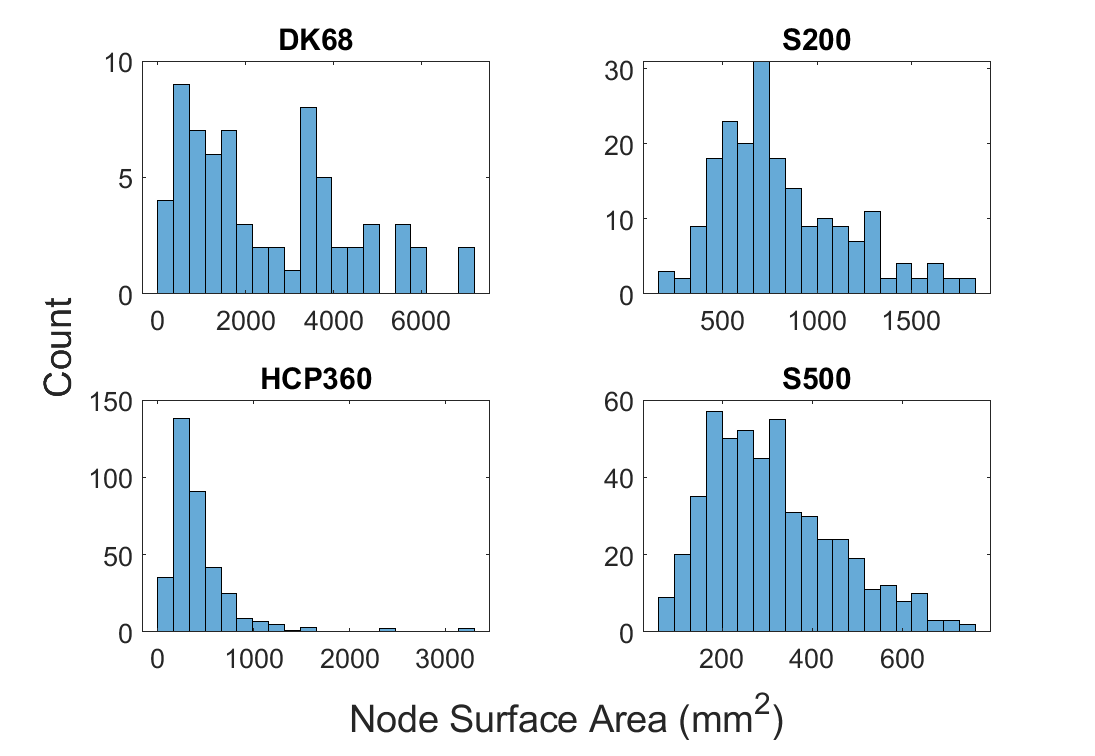

**Figure S15: Distributions of node surface areas in all parcellations**. The surface area for each node is given as the mean of the surface area for that node across participants. Note that the HCP parcellation is highly skew, and its largest node is larger than that of the S200 parcellation (3290 mm^2^ vs 1831 mm^2^).

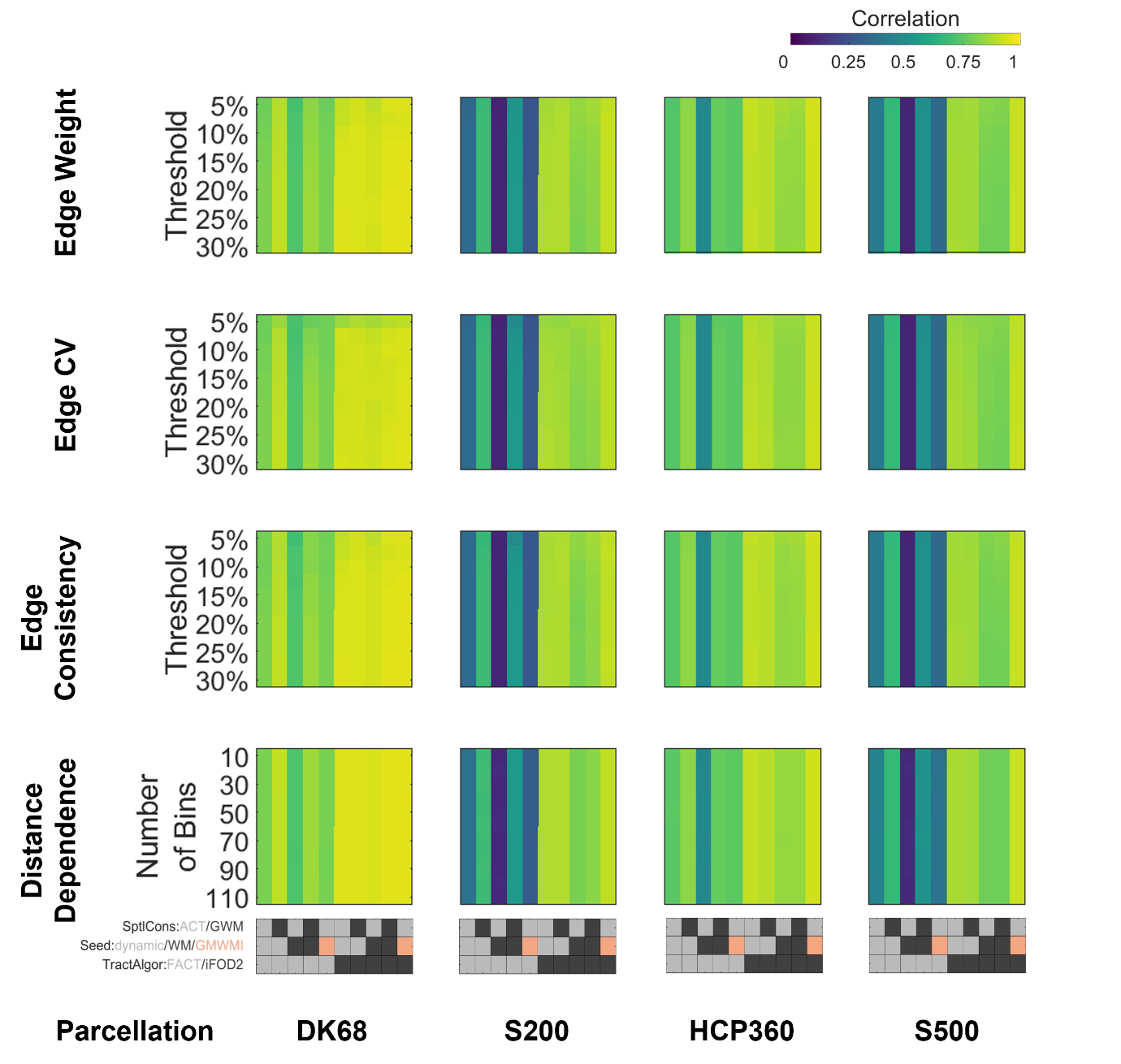

**Figure S16: Comparison of correlations between node strength and node surface area across all parcellations, tractography workflows, group reconstructions, and thresholds**. The parcellation and workflow vary along the x-axis, and the group reconstruction and threshold vary along the y-axis. Each subplot shows the correlation as a function of tractography workflow and threshold for a given parcellation and group reconstruction.

Parcellation: DK68 = Desikan-Killiany 68 nodes, S200 = Schaefer 200 nodes, HCP360 = Glasser 360 nodes, S500 = Schaefer 500 nodes. Tractography: SptlCons = spatial constraints on streamline propagation, ACT = anatomically constrained tractography, GWM = grey-white masking; Seed = streamline seeding algorithm, dynamic = dynamic seeding, WM = white matter seeding, GMWMI = grey matter-white matter interface seeding; TractAlgor = Streamline tractography algorithm, FACT = fiber assignment by continuous tractography, iFOD2 = second-order integration over fiber orientation distributions.

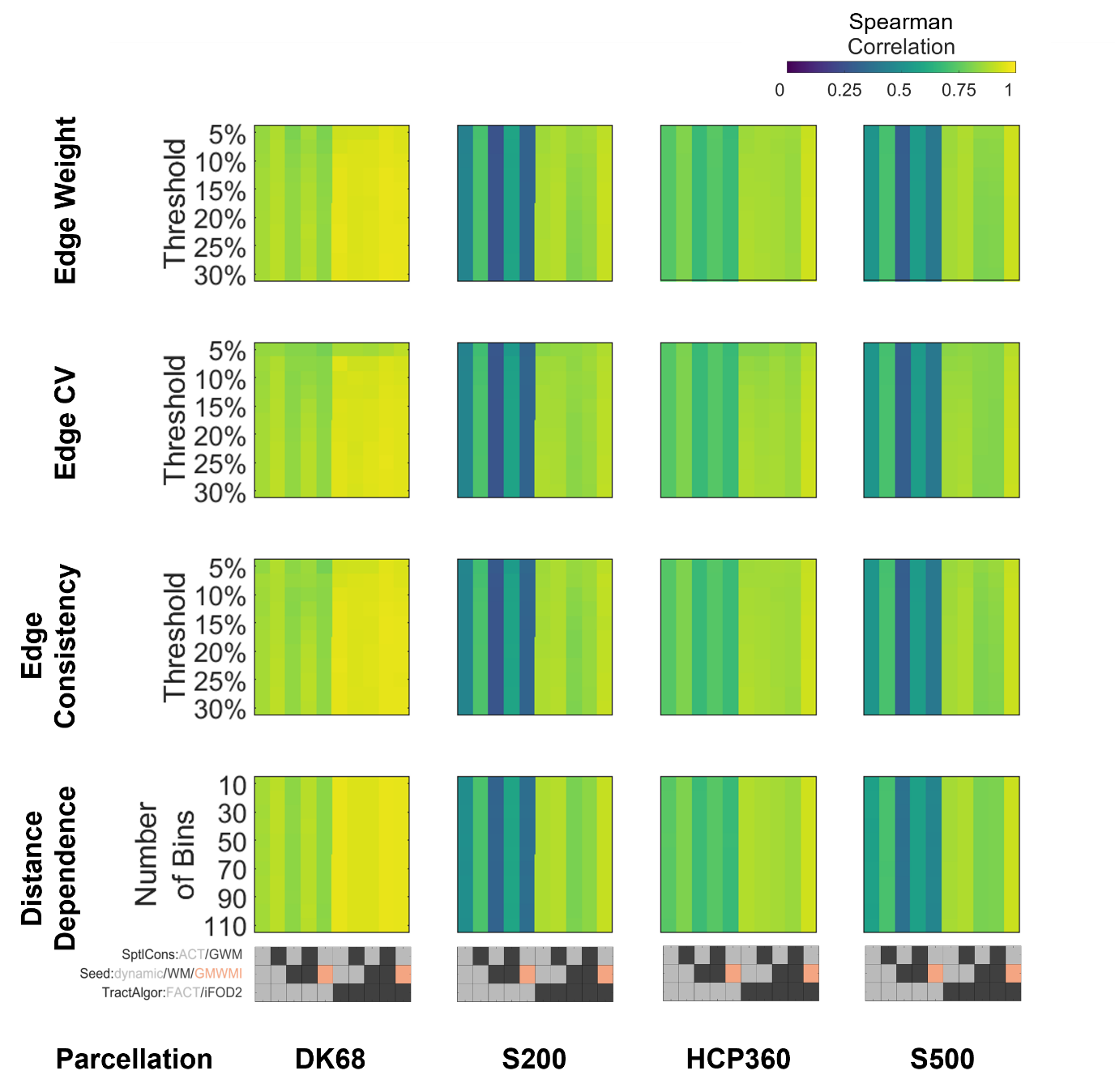

**Figure S17: Comparison of Spearman correlations between node strength and node surface area across all parcellations, tractography workflows, group reconstructions, and thresholds**. The parcellation and workflow vary along the x-axis, and the group reconstruction and threshold vary along the y-axis. Each subplot shows the correlation as a function of tractography workflow and threshold for a given parcellation and group reconstruction.

Parcellation: DK68 = Desikan-Killiany 68 nodes, S200 = Schaefer 200 nodes, HCP360 = Glasser 360 nodes, S500 = Schaefer 500 nodes. Tractography: SptlCons = spatial constraints on streamline propagation, ACT = anatomically constrained tractography, GWM = grey-white masking; Seed = streamline seeding algorithm, dynamic = dynamic seeding, WM = white matter seeding, GMWMI = grey matter-white matter interface seeding; TractAlgor = Streamline tractography algorithm, FACT = fiber assignment by continuous tractography, iFOD2 = second-order integration over fiber orientation distributions.

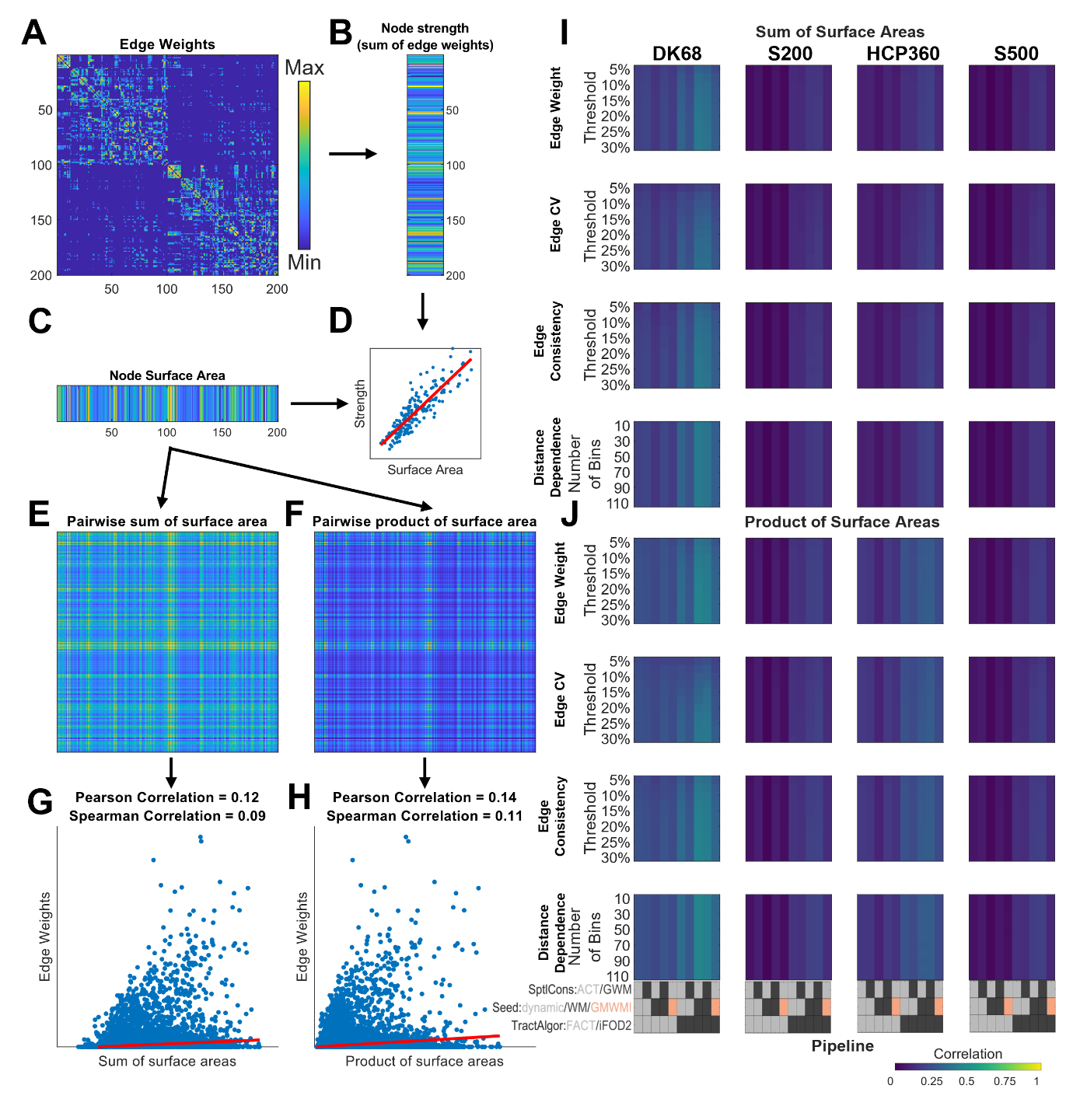

**Figure S18: Relationship between edge weight and endpoints’ surface area.** (A) Exemplar connectome (using the same parameters used in Figure 4A left; i.e., workflow 7) showing the weight of each edge. (B) Strength of each node; i.e.. sum of the edge weights. (C) Node surface area. (D) Relationship between node strength and node surface area (note that this figure is the same as Figure 4B left). (E, F) For each edge, its terminating nodes’ strengths are added/multiplied together. (G, H) Correlation between each edge’s weight (shown in panel A) and the sum/product of its nodes’ surface area (shown in panels E and F). (I, J) Correlation coefficient between edge weight and endpoints’ surface area (sum or product) across parcellations, workflows, and reconstructions.

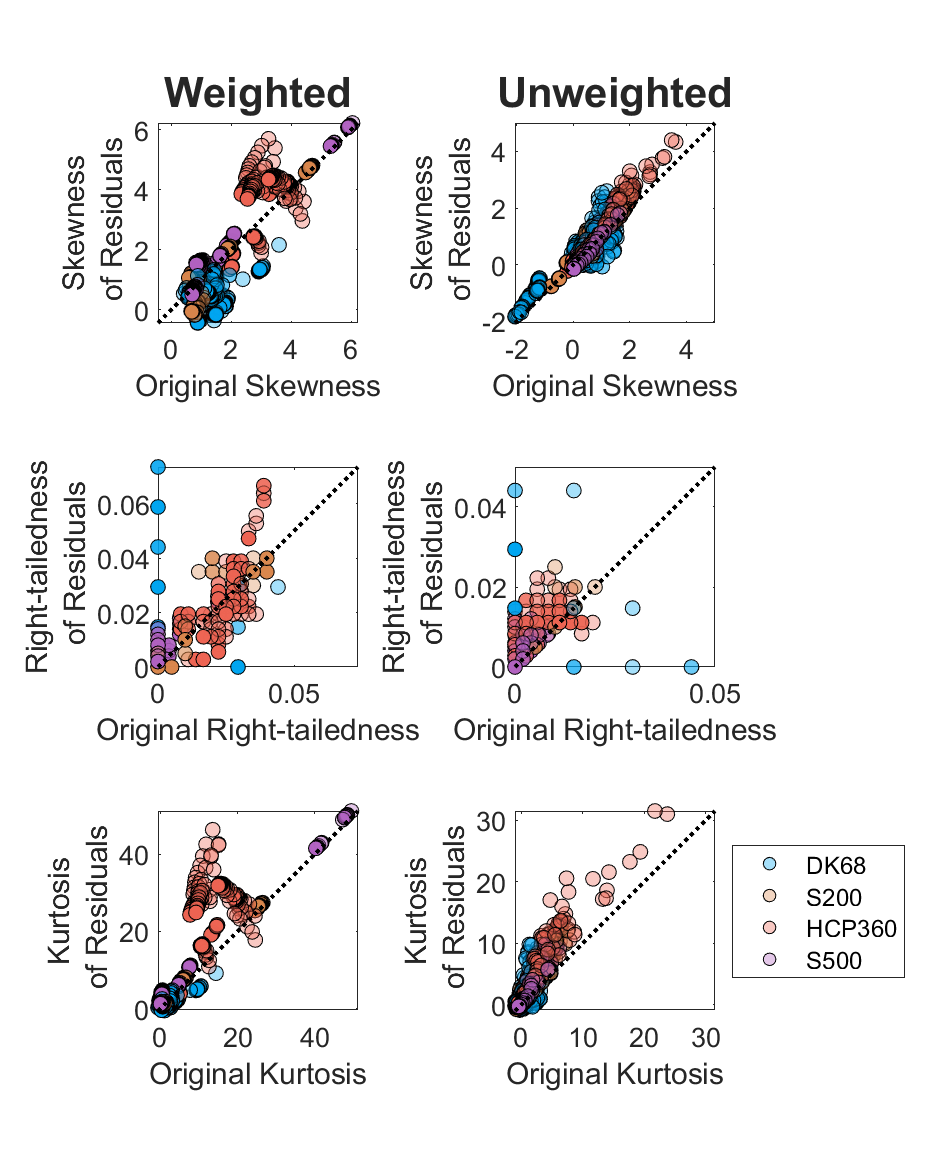

**Figure S19: Comparison of original and residual strength distributions**. Left, weighted connectomes; right, binarized connectomes. Top, skewness; middle, right-tailedness; bottom, kurtosis. All combinations of parcellations, tractography workflows, group reconstructions, and thresholds are shown. The lattice nature of the right-tailedness of the DK68 parcellation is due to the paucity of data points. The HCP360 parcellation has residuals that are slightly more skewed/kurtotic than the original but remains supra-exponential. Otherwise, the grouping of points along the line *y = x* suggests that the distribution of residuals is similar to the original strength distribution, at least until the fourth standardized moment.

.

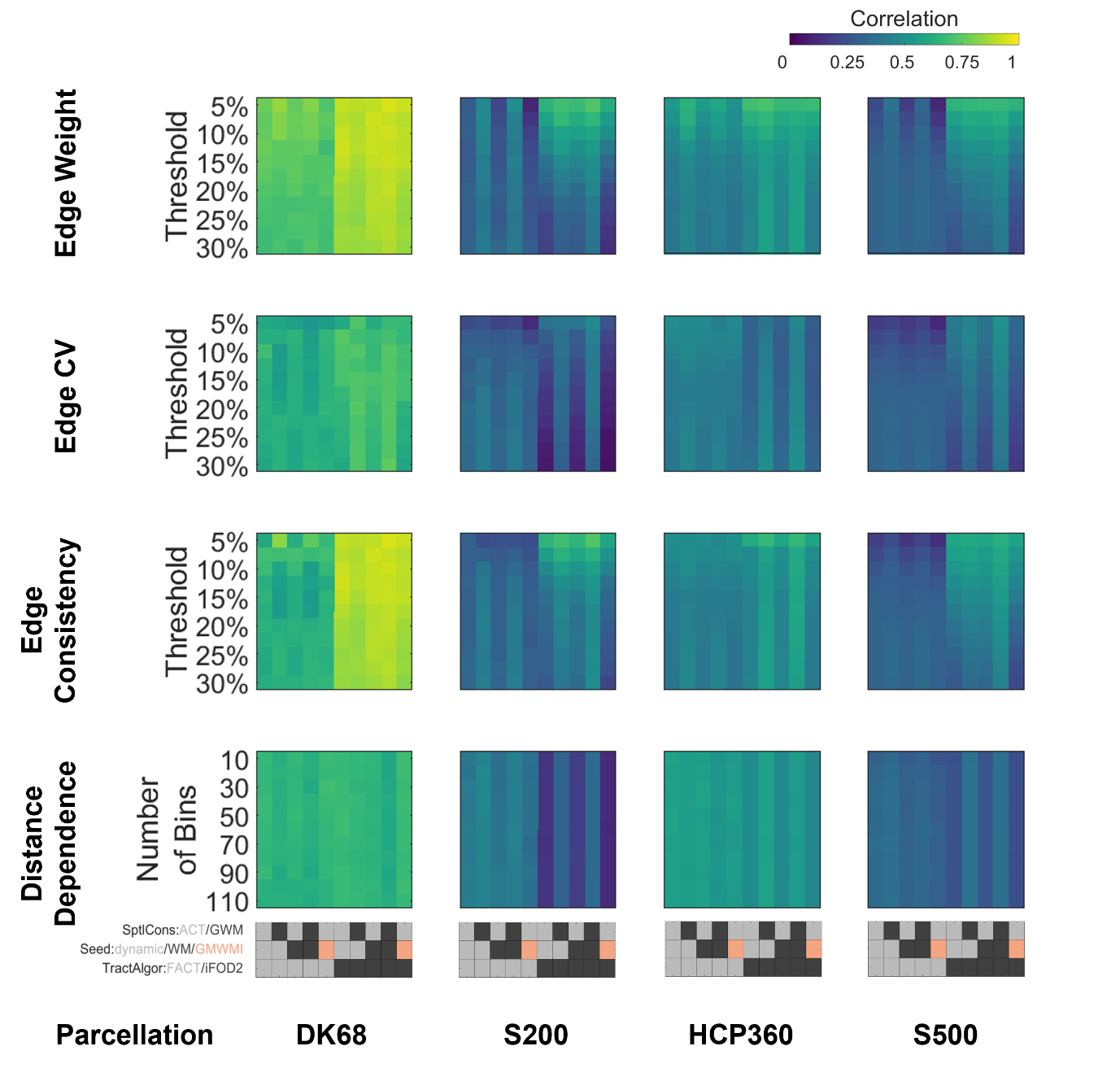

**Figure S20: Comparison of correlations between node degree and node surface area in binarized connectomes across all parcellations, tractography workflows, group reconstructions, and thresholds**. The parcellation and workflow vary along the x-axis, and the group reconstruction and threshold vary along the y-axis. Each subplot shows the correlation as a function of tractography workflow and threshold for a given parcellation and group reconstruction.

Parcellation: DK68 = Desikan-Killiany 68 nodes, S200 = Schaefer 200 nodes, HCP360 = Glasser 360 nodes, S500 = Schaefer 500 nodes. Tractography: SptlCons = spatial constraints on streamline propagation, ACT = anatomically constrained tractography, GWM = grey-white masking; Seed = streamline seeding algorithm, dynamic = dynamic seeding, WM = white matter seeding, GMWMI = grey matter-white matter interface seeding; TractAlgor = Streamline tractography algorithm, FACT = fiber assignment by continuous tractography, iFOD2 = second-order integration over fiber orientation distributions.
